# scBASE: A Bayesian mixture model for the analysis of allelic expression in single cells

**DOI:** 10.1101/383224

**Authors:** Kwangbom Choi, Narayanan Raghupathy, Gary A. Churchill

## Abstract

Allele-specific expression (ASE) at single-cell resolution is a critical tool for understanding the stochastic and dynamic features of gene expression. However, low read coverage and high biological variability present challenges for analyzing ASE. We propose a new method for ASE analysis from single cell RNA-Seq data that accurately classifies allelic expression states and improves estimation of allelic proportions by pooling information across cells.

---

Single-cell RNA sequencing (scRNA-Seq) can reveal features of cellular gene expression that cannot be observed in bulk RNA sequencing^1^. Allelic imbalance is common across many genes^2^ and can range from a subtle imbalance to complete monoallelic expression as in imprinted genes^3^ or genes under dosage compensation by X chromosome inactivation^4, 5^. Allele-specific expression (ASE) in single cells can provide a rich picture of the stochastic and dynamic properties of gene expression in individual cells. Analysis of single-cell ASE poses unique challenges due to the low depth of sequencing coverage per cell^6–13^. In addition, allelic proportions often form U-shaped or W-shaped distributions due to the occurrence of monoallelic transcriptional bursts.

We propose a novel method for the estimation of single-cell allele proportions, `scBASE`, in which we (1) disambiguate and count multi-mapping reads (multi-reads); (2) classify each gene in each cell into paternal monoallelic, bi-allelic, or maternal monoallelic expression states; and (3) address data sparsity by partial pooling of information across cells (Figure 1 and Methods). The *counting* step of `scBASE` applies an estimation-maximization (EM) algorithm to count multi-reads by *weighted allocation* to estimate expected read counts^14–16^. The *classification* and *estimation* steps are iterated and together achieve *partial pooling* of information among cells that are in the same allelic expression states. In the classification step, we compute the posterior probabilities of paternal, bi-allelic, and maternal expression states. In the estimation step we compute the posterior distributions of cell- and gene-specific allelic proportions. Read counting and partial pooling can be applied together, separately, or not at all. This leads to four different methods of estimating allelic proportions. In the unique reads methods (i), we estimate allelic proportions directly from the counts of uniquely mapping reads. In the weighted allocation method (ii), we apply the read counting step of `scBASE` to obtain estimated expected counts. We can apply the classification and estimation steps to either of these counts to obtain allelic proportions from unique reads with partial pooling (iii), or weighted allocation with partial pooling (iv). We have implemented these methods in extensible open-source software, `scBASE`, available at https://github.com/churchill-lab/scBASE.

**Figure 1:**
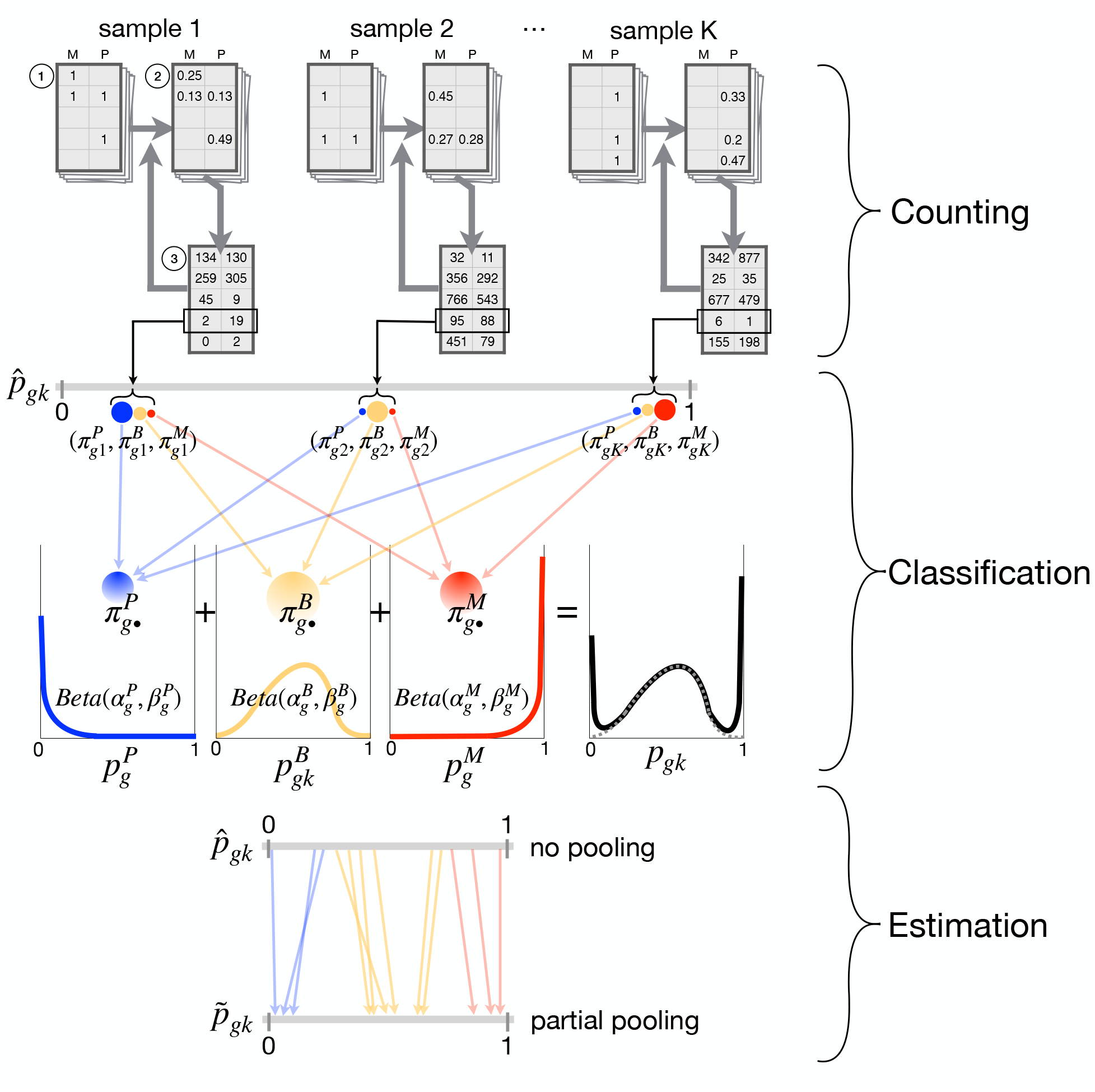
Overview of the scBASE algorithm. We summarize the three steps of the `scBASE` algorithm. The **Counting** step estimates the expected read counts using an EM algorithm to compute a weighted allocation of multi-reads. Each read is represented as an incidence matrix that summarizes all best-quality alignments to genes and alleles ①. Weighted allocation of multi-reads uses a current estimate of allele-specific gene expression to compute weights equal to the probability of each possible alignment ②. The weights are summed across reads to obtain the expected read counts for each gene and allele ③. Steps ② and ③ are repeated until the read counts converge. The weighted allocation estimates of maternal allelic proportion 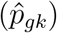 are obtained at this step. The **Classification** step computes the posterior probability of paternal monoallelic (P), bi-allelic (B), or maternal monoallelic (M) expression 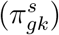 using current estimates of the model parameters (Equation 3 in Supplemental Methods). The classification model is a beta-binomial mixture model with three components. The model parameters are initialized to non-informative values and are obtained from the estimation step in subsequent iterations. The **Estimation** step uses to the classification results to re-estimate the weights of mixture components 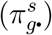 and parameters of the Beta densities 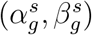 that define the distribution of the within-class the maternal allelic proportions 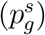. The partial pooling estimate of the maternal allelic proportions 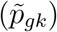 is obtained as an average of the class-specific proportions weighted by the class membership probabilities (Equation 1 in Methods).

In the following sections, we first examine the effects of weighted allocation on single-cell allele expression data. Then we evaluate the effects of partial pooling on estimation of allelic proportions. We then apply each of the four methods in `scBASE` to statistical testing of independence of allelic bursting. Finally, we illustrate the interpretive power of allelic expression by analysis of scRNA-Seq data from a development time course^7^.

## Results

We applied `scBASE` to scRNA-Seq data from 286 pre-implantation mouse embryo cells from an F1 hybrid mating between female CAST/EiJ (CAST) and male C57BL/6J (B6) mice^7^. Cells were sampled along a time course from the zygote and early 2-cell stages through the late blastocyst stage of development. We created a diploid transcriptome from CAST- and B6-specific sequences of each annotated transcript (Ensembl Release 78)^17^ and aligned reads from each cell to obtain allele-specific alignments. In order to ensure that genes had sufficient polymorphic sites for ASE analysis, we restrict attention to 13,032 genes that had at least 4 allelic unique reads in at least 10% of cells. Where indicated below, we apply `scBASE` to only 122 cells from the blastocyst stages of development, or to only 60 cells in the mid-blastocyst stage.

We first assessed the impact of weighted allocation of multi-reads on the estimation of allelic proportions. Any read that maps to one allele of one gene is a unique read. All other reads are multi-reads and they can be simple or complex. A read that maps uniquely to one gene but to both allelic copies is a simple allelic multi-read. A read that maps to multiple genes but only to one allele at each is a simple genomic multi-read. A read that maps to multiple genes as well as to both alleles in any of those genes are complex multi-reads. There are, in total, 9 patterns of simple and complex multi-read alignments for two genomic loci and two alleles (Supplemental Figure S1). We estimated unique reads and weighted allocation counts from each individual cell using all 286 cells to show how the number of monoallelic genes changes in each cell (Figure 2a). The sequence reads from these cells include 2.5% simple genomic multi-reads, 59.3% simple allelic multi-reads, and 23.3% complex multi-reads. In a typical scRNA-Seq workflow for ASE, these reads are discarded leaving only the unique 14.9% of the original sequence reads for analysis. This substantial loss of information could lead to high variability of allelic proportions and spurious findings of monoallelic gene expression. We find that using only uniquely mapping reads generates a higher rate of monoallelic expression calls (Figure 2a and Supplemental Figure S1), calling on average *∼*66 more genes with monoallelic expression in each cell. We also observed, on average, *∼*1,908 genes where the unique reads method fails to call bi-allelic expression compared to weighted allocation, for example, *Mtdh* (Figures 2b and 2c). These genes are consistently bi-allelic in many cells according to weighted allocation, but their pattern of allelic expression based on unique reads can be misinterpreted as monoallelic expression and, as a result, allelic bursting appears to be more dynamic.

**Figure 2:**
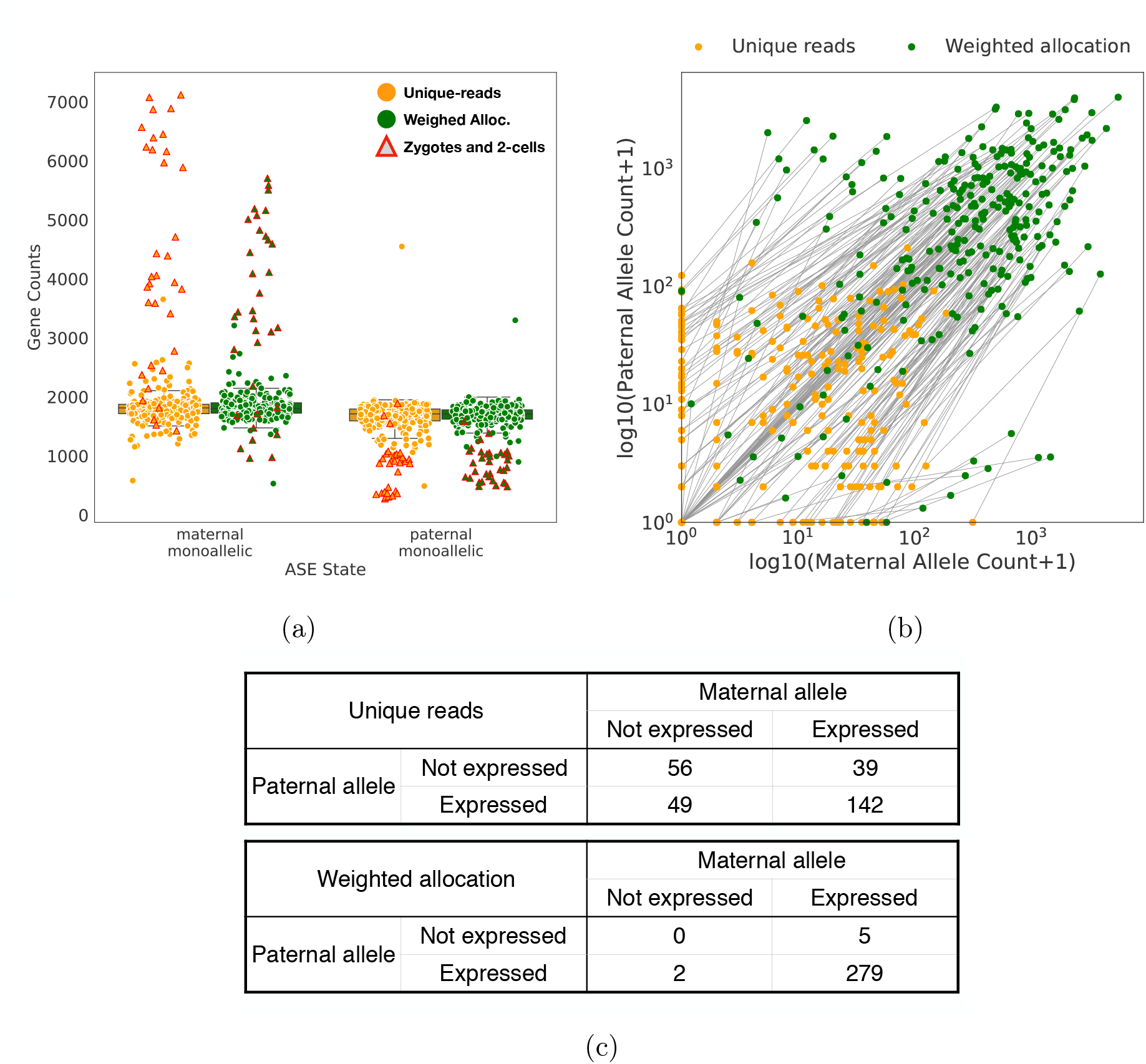
Weighted allocation of multi-reads reduces monoallelic expression calls. (a) For each of 13,032 genes, we obtained the allele-specific read counts by unique reads and by weighted allocation. We counted the numbers of genes in each cell that showed either maternal or paternal monoallelic expression and display the results as points (one per cell) overlaid on boxplots. Each data point in this figure represents a cell and we are showing all 286 cells including zygote and 2-cells (highlighted in red). The zygote and 2-cell stage cells have large numbers of genes with maternal monoallelic expression. On average there are 66 fewer monoallelic calls per cell with the weighted allocation counts. The outlier cell with high levels of paternal monoallelic expression was noted in Deng et al.^7^. **(b)** We selected one gene (*Mtdh*) to illustrate the distribution of maternal (X-axis) and paternal (Y-axis) counts across 286 cells. The weighted allocation counts (green) are connected to their corresponding unique counts by a line in the scatter plot. **(c)** Cross-tabulation (2 2 table) of maternal and paternal allelic expression for *Mtdh* gene with unique reads and weighted allocation counts. The unique counts resulted in 88 cells with monoallelic expression while only 7 monoallelic calls were seen with weighted allocation.

Next we evaluated the impact of partial pooling on the estimation of allelic proportions. Since it is best to apply partial pooling to each cell type separately, we focus attention on the 122 mature blastocyst cells, the largest group in Deng et al.^7^ data. These cells have the coverage of *∼*14.8M reads per cell in average, and we down-sampled these data by randomly selecting 1% of reads to obtain an average coverage of *∼*148k reads per cell. We estimated allelic proportions using each of four methods: (i) unique reads, (ii) weighted allocation, (iii) unique reads with partial pooling, and (iv) weighted allocation with partial pooling.

We compared the estimated allelic proportions from the down-sampled data to estimates obtained from the full data using the corresponding unique reads or weighted allocation estimates with no pooling. The full data are based on 100-fold more reads per sample and provide an approximate truth standard. A similar approach to evaluation of single-cell data analysis was employed by Huang et al.^18^. In order to assess the effects of partial pooling, we computed differences in the mean squared error (MSE) of estimated allelic proportions with and without partial pooling. Partial pooling applied to the unique read counts improves estimation for the majority of genes (4,392 versus 1,367 out of 5,759 genes) with an average MSE difference of 0.018 (Figure 3a). Partial pooling applied to the weighted allocation counts improves estimation for most genes (5,078 versus 1,673 out of 6,751 genes) with an average MSE difference of 0.016 (Figure 3b). In both cases, the greatest gains are seen in the low expression range (<10 unique reads per gene). For the most highly expressed genes, there is little or no reduction in MSE, which is consistent with our expectation that pooling of information across cells is most impactful when coverage is low.

**Figure 3:**
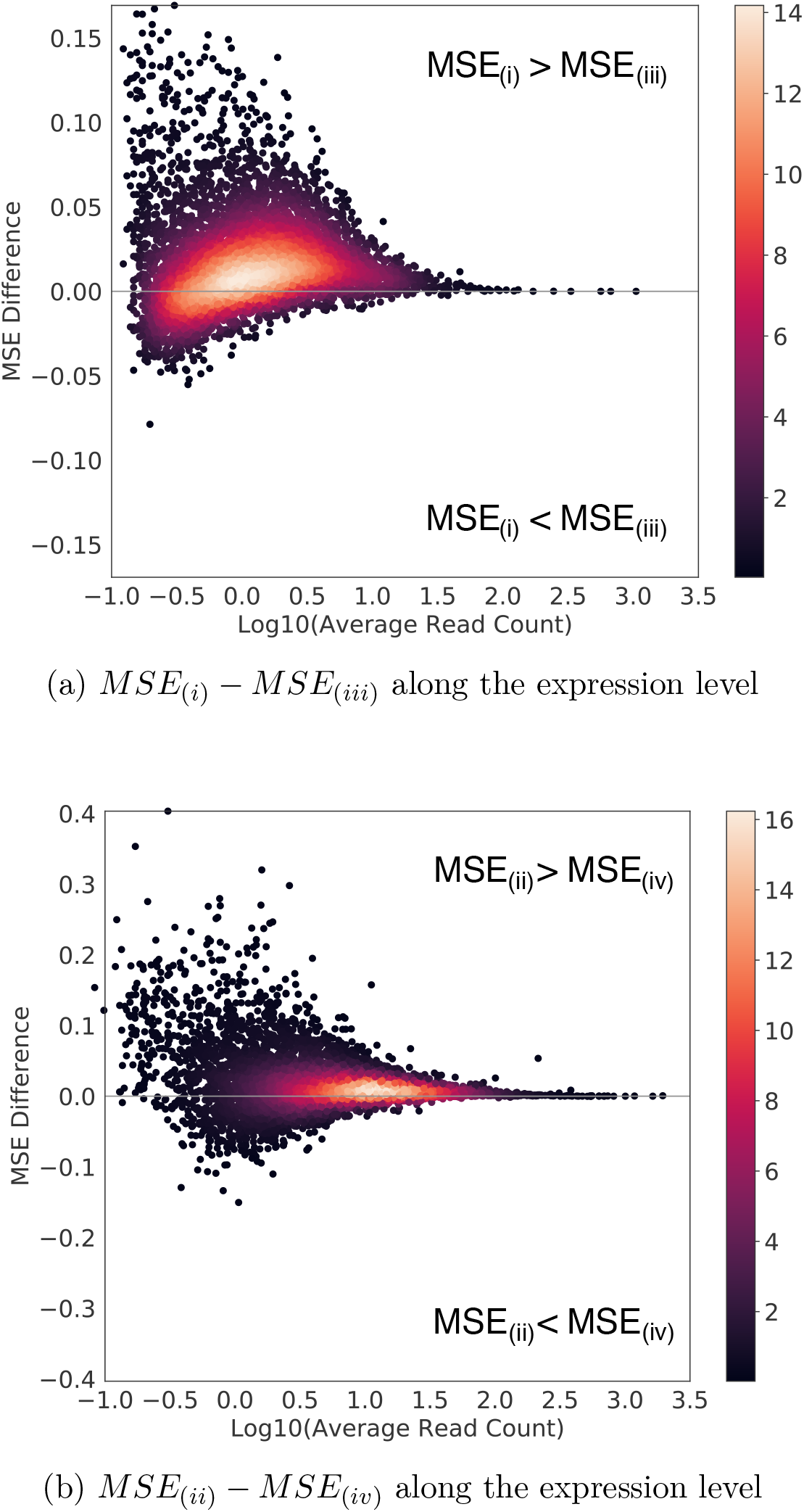
Partial pooling improves the accuracy of estimated allelic proportions. We randomly sampled 1% of reads from the full data of 122 mature blastocyst cells to obtain a sub-sample of 147,538 reads per cell, on average. We estimated gene- and cell-specific allelic proportions from the sub-sampled data, and computed mean squared error (MSE) between the estimated allelic proportions from the full data versus the sub-sampled data. We compared the MSE based on partial pooling versus the MSE from no pooling estimates, and display the difference on the y-axis along the expression level in unique-read counts on the x-axis. We made this comparison for **(a)** unique reads and for **(b)** weighted allocation. Points representing individual genes are shown as a density heatmap.

The timing of allelic bursting events is a defining feature of stochasticity in gene expression^19^. One fundamental question is whether the occurrence of allelic bursts is coordinated or if bursts occur independently for each allele. Statistical independence of maternal and paternal bursting can be evaluated using a 2*×*2 table of counts of the numbers of cells for which a given gene is expressed only from the maternal allele, only from the paternal allele, from both, or not expressed (as in Figure 2c). If allelic bursts occur independently, the log-odds ratio (logOR) computed from this 2 *×* 2 table should be close to zero. In order to relate this standard approach^20^ for testing the independence hypothesis to alternative methods^7, 21^ that have been proposed for scRNA-Seq data, it is helpful to consider a geometric representation of the proportions of cells in each allelic expression state (Figure 4a). Proportions are numbers greater than or equal to zero that sum to one. They can be represented as a point in a 3-dimensional tetrahedron in 4-dimensional space – the 4D simplex^22^. When maternal and paternal bursting events occur independently, the proportions should fall near the 2-dimensional surface within the simplex where the logOR is equal to zero (cross-hatched region in Figure 4a). The method of testing independence used in Deng et al.^7^ and Larsson et al.^21^ imposes an additional constraint on the 2 *×* 2 table proportions by assuming that the frequencies of maternal and paternal bursting events are equal (*p*_*M*_ = *p*_*P*_). This constraint corresponds to a 2-dimensional cross-section of the simplex, indicated by the blue triangle in Figure 4a. Projection of points in the 4D simplex onto this triangle produces the graphical representation used by Deng et al. (e.g., Figure 4b). This illustrates how the Deng et al. method is a special case of the logOR test.

**Figure 4:**
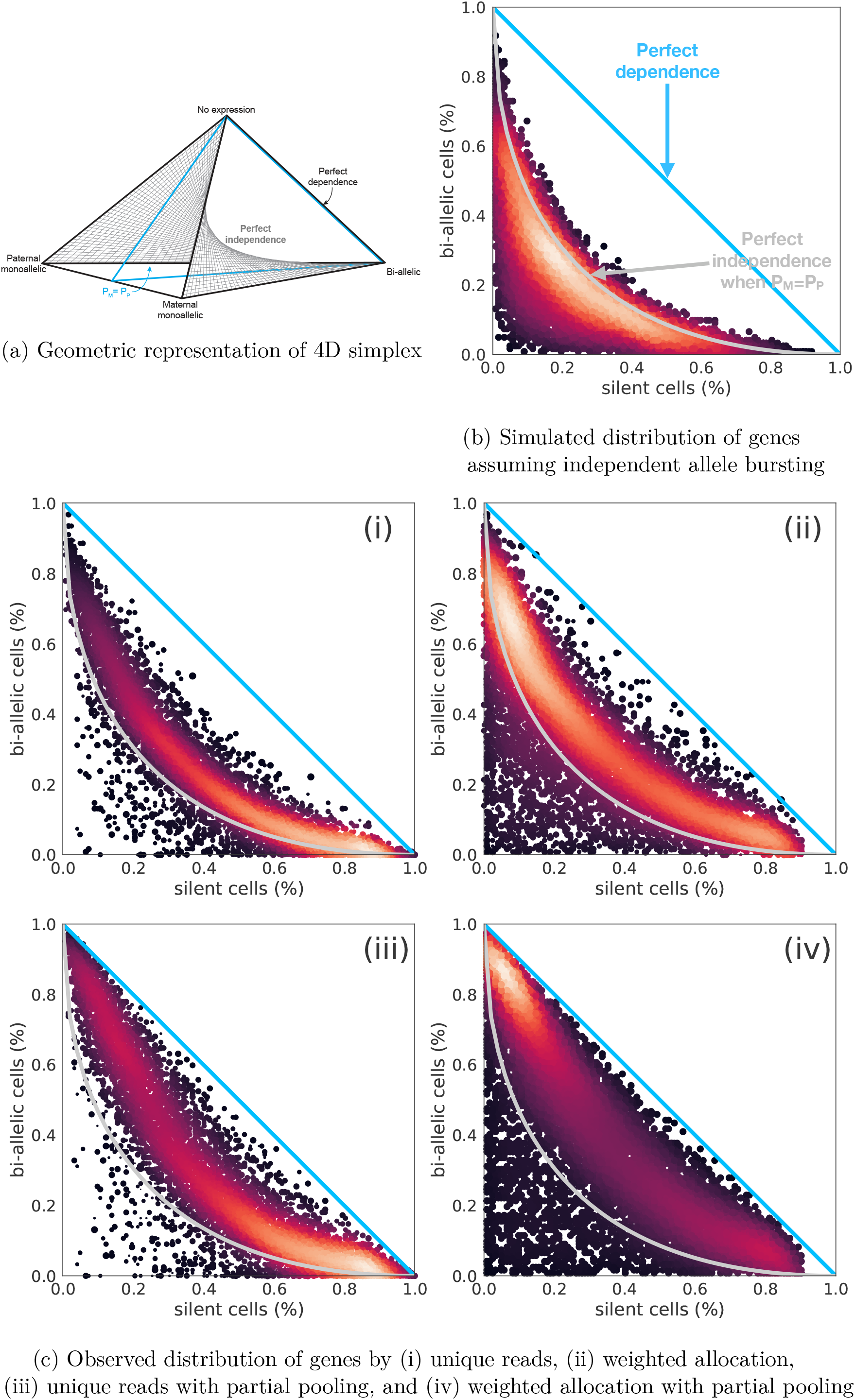
Independence of allelic bursting. **(a)** The geometry of the 2×2 table proportions can be represented as a simplex, a 3D tetrahedral region of 4D space in which proportions are all non-negative and sum to one. The vertices of the simplex correspond to genes where all cells are in the same allelic expression state as indicated by labels. The distance from a vertex is inversely related (1-x) to the proportion of cells in that state. The shaded surface inside the simplex represents proportions corresponding to the perfect independence model, i.e., the logOR equals zero. The blue triangle indicates proportions with equal maternal and paternal expression *p*_*M*_ = *p*_*P*_. **(b)** We simulated data under the perfect independence model without assuming *p*_*M*_ = *p*_*P*_ and plotted the proportions of bi-allelic and silent cells as in Deng et al.^7^. **(c)** Four panels illustrate the proportions of bi-allelic and silent cells as estimated from (i) unique reads, (ii) weighted allocation, (iii) unique reads with partial pooling, and (iv) weighted allocation with partial pooling. Points representing individual genes are shown as a density heat map.

We evaluated bursting independence on the 122 mature blastocyst cells as was done in Jiang et al.^23^. We first simulated data under the assumption of independent allelic bursting (Methods) and plotted the results to illustrate how points will be distributed in this diagram when the pure independence model is true with and without the constraint of *p*_*M*_ = *p*_*P*_ (Figure 4b). Next we estimated the 2 *×* 2 table proportions of allelic expression states using each of the four methods (i*∼*iv) implemented in `scBASE`. The appearance of the data in Figures 4c is qualitatively distinct from the simulated data (Figure 4b). Moreover, the null hypothesis of independence is rejected by the method used in Jiang et al.^23^ for the majority of genes regardless of the method used to estimate the allelic state proportions (Supplemental Figure S2a). `SCALE` reports 3,381 genes that are non-independent using the results of unique-reads method, 4,815 genes using weighted allocation, 6,068 genes by unique reads with partial pooling, and 6,761 genes based on weighted allocation with partial pooling at the FDR level of 5%. Similarly the logOR are away from 0 for thousands of genes. For example, 2,845 and 3,763 out of 8,290 genes had |logOR|>2 using unique reads and weighted allocation. More genes have |logOR|>2 after partial pooling with `scBASE`: 5,622 and 6,209 respectively. The majority of genes had positive logOR, indicating a tendency for bursting to occur more in synchrony than chance would predict (Supplemental Figure S2b). We repeated this analysis using three additional data sets^21, 24, 25^ and arrive at similar conclusions in each case (Supplemental Figures S3, S4, and S5). The evidence for statistical dependence of bursting is strong and application of weighted allocation and partial pooling strengthens this conclusion.

The `scBASE` classification step provides a novel way to characterize allelic imbalance across a population of cells by estimating the expected proportions of cells in different transcriptional states. Using `scBASE`, we can compute the posterior probability of allelic expression states of genes in each cell. This probabilistic classification allows for uncertainty associated with statistical sampling from the pool of transcripts that are present in the cell including the occurrence of zero read counts. Based on the posterior probabilities, we can derive the expected proportions of cells in states P, B, and M, which can be represented as point in a triangular simplex diagram. (Note that this representation is a projection of points in the 4D simplex onto the bottom triangular region, Figure 4a.) The classification step of `scBASE` assumes that all genes are expressed at some level, which may be very low for some genes. This allows us to classify the allelic expression of cells that may have zero read counts due to statistical sampling. To interpret the distribution of allelic expression across cells, we designate seven patterns of allelic expression (Figure 5a). Genes that are predominantly expressed as P, B, or M will appear near the corresponding vertex of the triangle (**P**, **B** or **M** region). Genes with mixed allelic states will appear along the edges (**PB**, **BM**, or **MP** region) or near the center of the triangle (all three states, **PBM** region). For example, the gene *Pacs2*, which is expressed from either the maternal or the paternal allele but rarely both, is classified as an **MP** gene. The bi-allelic region (**B**) includes genes that are consistently expressed from both alleles e.g., *Mtdh*. The **PB** and **BM** regions include genes that show a mixture of bi-allelic and monoallelic expression with a strong allelic imbalance, e.g., *Timm23* and *Tulp3*. The majority of genes (56.9%) in the blastocyst stages of development are in the **PBM** region (Supplemental Figure S6). These genes display a mix of mono- and bi-allelic expression states (e.g., *Akr1b3*) that is consistent with dynamic allele-specific gene expression with a low bursting rate relative to mRNA half life.

**Figure 5:**
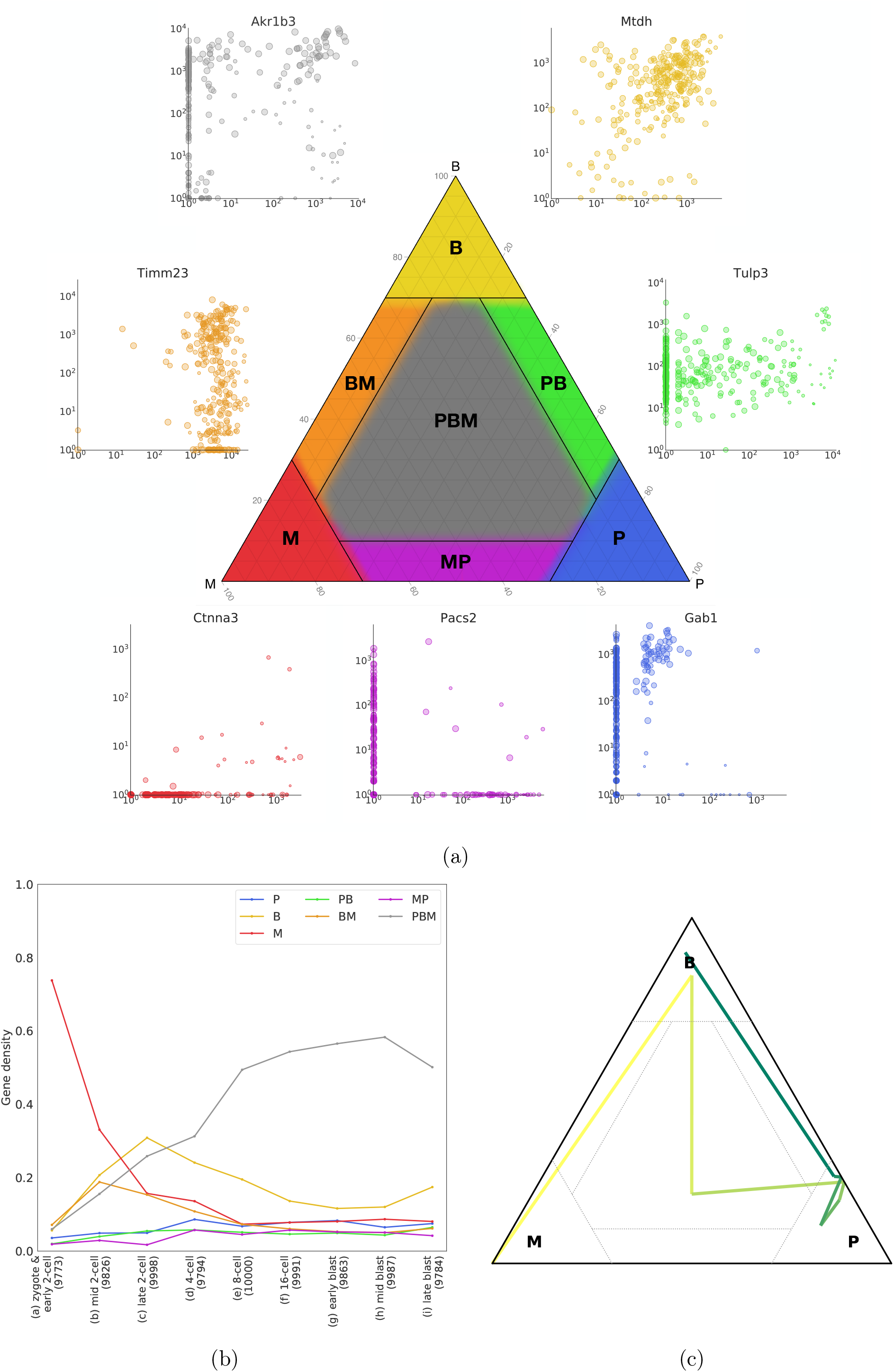
Classification of allele-specific expression patterns across cells. **(a)** For each gene in each cell, the classification step of scBASE estimates allelic state probabilities 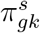, where *s* indicates paternal monoallelic (P), bi-allelic (B), or maternal monoallelic (M) expression. The average proportions of cells in each allelic state 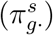 can be represented as a point in a triangular diagram which is a 3D simplex corresponding to the projection of points onto the bottom triangular region of the 4D simplex in Figure 4a. A gene that is predominantly paternal, bi-allelic, or maternal across the cell population will be plotted near the corresponding vertex. Points representing genes with mixed classification states across the cell population will appear along the edges or in the center of the triangle. We delineate seven patterns of allelic expression for a gene as indicated by the different colored regions in the diagram: **P** (blue), **B** (yellow), **M** (red), **PB** (green), **BM** (orange), **MP** (purple), and **PBM** (gray). Examples of genes from each pattern are shown as scatter plots of maternal and paternal read counts (log10 scale). Each point in the scatter plot corresponds to one cell (n=286 embryo cells). For example, the gene *Pacs2* is expressed from either the maternal or the paternal allele but rarely both and is classified as an **MP** gene. The bi-allelic region (**B**) includes genes that may show allelic imbalance 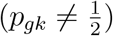 across many cells but consistently express both alleles (e.g., *Mtdh*). The **PB** and **BM** regions will include genes that show a mixture of bi-allelic expression and monoallelic expression. Many of the genes in these regions have strong allelic imbalance and cells with monoallelic expression could be due to statistical sampling zeros in the lower expressed allele (e.g., *Tmim23* and *Tulp3*). The expression pattern in blastocyst cells for the majority of genes (57%) fall in the **PBM** region and display a pattern that is a mix of mono- and bi-allelic expression states across cells (e.g., *Akr1b3*). **(b)** Cells were divided into nine developmental stages as indicated on the X-axis. The cell types and numbers of expressed genes at each stage are indicated in parentheses on the X-axis. For each stage, we counted the proportion of expressed genes that fall into each of the seven allelic expression patterns (Y-axis), indicated by lines using the same color coding used in Figure 5a. In the zygote and early 2-cell stage, most genes show purely maternal expression (**M**). The proportion of maternally expressed genes decreases through subsequent stages of development. The numbers of genes showing purely paternal expression (**P**) is low across all developmental stages. The **M** and **P** classes become equally represented in the later stages of development. The 2- and 4-cell stages show high levels of bi-allelic expression (**B**) and the mixed class (**PBM**) proportion becomes highest by the 8-cell stage. **(c)** The expected proportions of cells in each allelic state 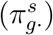 for one gene *Akr1b3* at each stage of the developmental time course is shown as a trajectory in the 3D simplex. Yellow to blue color line segments indicates the transitions between developmental stages. This gene starts in the maternal monoallelic state (**M**), it transitions through **PBM** to a paternal expression state (**P**), and then transitions to bi-allelic expression (**B**) in the blastocyst stages.

We applied `scBASE` (with weighted allocation and partial pooling) to track changes in the ASE patterns of cells sampled over a developmental time course (Figure 5b, Supplemental Figure S6 and S7). Our aim is to classify allelic state distributions within subpopulations of cells defined by developmental stages. To achieve this, we first ran `scBASE` MCMC algorithm on all 286 cells to estimate the prior parameters, 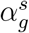 and 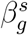(Figure 1 and Supplemental Methods). These parameters describe the distribution of allelic proportions in each allelic state. According to our diagnostic criteria, `scBASE` MCMC algorithm produced reliable parameter estimation for 10,017 out of 13,032 genes. We then ran `scBASE` EM algorithm (with the prior parameters fixed) on each subpopulation of cells to estimate developmental stage-specific parameters (Details are provided in Methods.). In the zygote and early 2-cell stages, essentially all genes show monoallelic maternal expression. At this stage, the hybrid embryo genome is not being transcribed and the mRNA present is derived from the mother (inbred CAST genome). At the mid 2-cell stage the hybrid embryo is being transcribed and we start to see expression of the paternal allele for some genes. Many genes exhibit the **M** and **BM** patterns through the 8- or 16-cell stages perhaps due to the persistence of long-lived mRNA species that were present at the 2-cell stage. The bi-allelic class **B** dominates the late 2-cell and 4-cell stages indicating high levels of expression at rates that exceed the half-life of most mRNA species. In the later stages of development, 8-cell through late blastocyst, most genes transition into the **PBM** pattern.

There are *∼*400 genes that make dramatic transitions across allelic expression states. For example, *Akr1b3* (Figure 5c) starts in the zygote and early 2-cell stage with only maternal alleles present. It transitions to bi-allelic expression by the mid 2-cell stage indicating the onset of transcription of the paternal allele. It then transitions through the paternal monoallelic state. Our interpretation is that the early maternally derived transcripts were present prior to fertilization and these transcripts are still present when the paternal allele in the hybrid embryo gene starts to express. The early maternal transcripts are largely degraded by the 4- to 8-cell stages where we see only expression from the paternal allele. In the early blastocyst stages, we start to see embryonic expression of maternal alleles resulting in a bi-allelic expression pattern by the late blastocyst stage.

## Discussion

Allelic expression in single cells has provided new insights into the dynamic regulation of gene expression^24^. However, estimates of allelic proportions can display high statistical variation due to low depth of sequencing coverage per cell. The common practice of discarding multi-mapping reads exacerbates this problem. The `scBASE` algorithm reduces statistical variability by retaining and disambiguating multi-read data. It further improves estimation of allelic proportions by partial pooling of information across cells in the same ASE states. As a result we can obtain a more precise and accurate picture of gene expression dynamics in which biological stochasticity is revealed by reducing statistical variation.

Weighted allocation has been demonstrated to improve gene expression estimation in whole-tissue RNA-Seq^14–16^. When estimating total gene expression with weighted allocation, only genomic multi-reads need to be resolved and these typically represent a small proportion of all reads. When estimating allele-specific expression, however, depending on the levels of nucleotide heterozygosity, the majority of reads may lack distinguishing polymorphisms and will be allelic multi-reads. Complex multi-reads with ambiguity in both genomic and allelic alignment can carry useful information about allele-specific expression, as illustrated in Supplemental Figure S1.

`scBASE` uses partial pooling in the context of a mixture model with three allelic expression states (paternal monoallelic, bi-allelic, and maternal monoallelic) to preserve cell-to-cell heterogeneity by pooling information across cells that are in the same state. Combining information across cells, therefore, does not weaken the signals of strong allelic imbalance. We applied `scBASE` to X chromosome genes in female cells of three different data sets^7, 24, 25^. In the Reinius et al. fibroblast data, partial pooling corrected the allelic proportions of Xist gene expression towards either maternal or paternal monoallelic expression for both unique reads and weighted allocation counts (Supplemental Figure S9a). Looking at expression of all X chromosome genes in these same cells, we observe that partial pooling strengthens the expected pattern of expression due to X chromosome inactivation (XCI) consistent with Xist allele expression (Supplemental Figure S9b). We observe that XCI is often incomplete and not uniform across cells. In the Chen et al. and Deng et al. data sets, Xist is clearly in the bi-allelic expression state in many of mouse embryo cells, epistem cells, or motor neuron cells and this is preserved after partial pooling. We also observe that XCI is not fully established for these cells (Supplemental Figure S10, and S11). In addition, for genes that are reported to be imprinted^26–28^ we examined their allelic expression. Irrespective of the estimation method applied, many of these genes do not appear to be fully imprinted in these three data sets (Supplementary Figure S12 and S13). However, for those genes that do show evidence of imprinting, i.e., appear in **M**- or **P**-class, partial pooling improves the evidence for monoallelic expression for both unique reads and weighted allocation counts.

The `scBASE` analysis incorporates statistical uncertainty in both the classification of allelic expression state and the estimated allelic proportions of a gene. To evaluate the precision of the estimated parameters, we have computed the posterior standard deviation of allele proportions across a range of total read counts and with varying numbers of cells (286 cells versus 60 cells). The trends are as expected, deeper read coverage or more cells improves the precision of estimation (Supplemental Figure S8). Our probabilistic classification accounts for uncertainty and can estimate the allelic expression state of a gene even when few or no reads are sampled from a given cell based on the behavior of other cells. The `scBASE` model is still reliable with degenerate inputs, for example, in the most extreme case of a single cell and a gene with zero total reads, the algorithm provides a sensible answer: class probabilities are 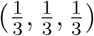 and a nearly uniform distribution for allelic proportion (mean at 0.5 with standard deviation of 0.2), indicating that the data does not contain any information. As the number of cells or the read depth increases, the class probabilities become more concentrated and the posterior distribution for the allelic proportion gets narrower. Partial pooling has the biggest impact when read coverage is low and the number of cells is large (Figure 3 and Supplemental Figure S8).

`scBASE` software can be implemented as part of a scRNA-Seq analysis pipeline. For example, we applied `SCALE` software using counts based on four methods: (i) unique reads, (ii) weighted allocation, (iii) unique reads with partial pooling, and (iv) weighted allocation with partial pooling implemented in `scBASE`. We found that more genes appeared to be non-independent when weighted allocation-based counts are used in `SCALE`. Even more genes were identified as non-independent using counts based on partial pooling (Results and Supplemental Figure S2a). Although it is not mentioned in Jiang et al.^23^, a substantial number (3,485 at FDR=5%) of genes were identified as non-independent using the allelic counts (unique reads) reported by Deng et al. Our findings suggest that running `SCALE` with `scBASE` estimated read counts as input will result in more accurate estimates of bursting kinetics and reduced levels of monoallelic gene expression when compared to results obtained using unique read counts.

The statistical properties of allelic bursting shed light on the nature of gene expression regulation. If expression bursts are statistically independent, this would imply that the regulation of allelic expression is local and acting autonomously at each allele. Under the perfect independence model, there would be no shared regulation of expression across alleles and the counts of cells in each allelic state will satisfy statistical criteria for independence. Under an alternative model, perfect dependence, bursting would be precisely coordinated across alleles and bursts would occur synchronously. All cells would be in either the bi-allelic or not expressed states. Our analysis of published scRNA-Seq data from four different experiments^7, 21, 24, 25^ indicates that neither of these extremes is true (Figure 4 and Supplemental Figure S2, S3, S4, and S5). We observed that the pattern of bursting is statistically dependent and positively correlated (logOR > 0) for the majority of genes. It is neither statistically independent nor perfectly synchronous. This suggests that regulation of allelic expression has both shared and locally autonomous components. While our statistical analysis cannot identify the mechanisms of regulation, it seems plausible that diffusible transcription factors could be responsible for the coordinated component of regulation. Local control is likely to be cis-acting and may involve stochastic variation in the activation of the transcriptional machinery. Additional experimental work would be required to test these hypotheses and to identify the cis-acting molecular events that trigger bursting of gene expression. However, the available data are sufficient to reject both hypotheses of perfect independence and of perfect dependence of allelic bursting.

When estimating parameters associated with many genomic features in each of many individual cells, one can improve the estimated parameters by pooling information across cells. The motivation behind partial pooling is that the individual estimates are unbiased but lack precision whereas the average provides a precise but biased estimate for individual cells and also masks cell to cell heterogeneity entirely. Weighted allocation of multi-mapping reads is not just to avoid information loss but is effective to prevent possible bias due to the genomic multi-reads that contain allele information. For these reasons, we generally recommend the strategy (iv) weighted allocation with partial pooling. But we provide all four options in `scBASE` so more evaluation could be performed in other contexts. These general principles –– retention of multi-mapping sequence reads and partial pooling of information across cells –– apply broadly to analysis of genomic sequencing data but they are especially critical in single cell applications where the observed numbers of reads for each gene in each cell may be very small.

## Methods

### Data

Deng et al.^7^ sampled 286 pre-implantation embryo cells from an F1 hybrid of CAST*×*B6 along the stages of prenatal development. Embryos were manually dissociated into single cells using Invitrogen TrypLE and single-end RNA-Seq sequencing was performed using Illumina HiSeq 2000 (Platform GPL12112). We downloaded the data, Series GSE45719 from Gene Expression Omnibus (GEO) at http://www.ncbi.nlm.nih.gov/geo/query/acc.cgi?acc=GSE45719. There were fastq-format read files for 4 single-cell samples from zygote stage, 8 from early 2-cell, 12 from mid 2-cell, 10 from late 2-cell, 14 from 4-cell, 47 from 8-cell, 30 from 16-cell, 43 from early blastocyst, 60 from mid blastocyst, and 58 from late blastocyst stage. The Reinius et al. data^24^ consist of primary mouse fibroblast cells from the F1 reciprocal crosses of CAST*×*B6 (125 cells, sex-typed) and B6*×*CAST (113 cells, sex-typed), available from GEO at https://www.ncbi.nlm.nih.gov/geo/query/acc.cgi?acc=GSE75659. The Chen et al. data^25^ are from mouse embryonic stem cells (mESCs) from an F1 hybrid of B6*×*CAST: 111 mESCs cultured in 2i and LIF, 120 mESCs cultured in serum and LIF, 183 mouse Epistem cells (mEpiSCs), and 74 post-mitotic neuron cells. The samples are sex-typed. We downloaded SRA format files available at https://www.ncbi.nlm.nih.gov/geo/query/acc.cgi?acc=GSE74155. Larsson et al.^21^ generated 224 individual primary mouse fibroblast cells from the F1 hybrid of CAST*×*B6. As the data are from non-standard SMART-Seq2 platform, we downloaded the allele-specific UMI counts from https://github.com/sandberg-lab/txburst/tree/master/data (as of April 19th, 2019), and we were unable to apply weighted allocation to these data.

### scRNA-Seq read alignment

For the F1 hybrid mouse we aligned reads to a phase-known diploid transcriptome – this is a best-case scenario for phasing. When dealing with more complex genomes, phasing should be performed beforehand if haplotype-specific transcriptomes are not available and scphaser^29^ is one possible approach. We reconstructed the CAST genome by incorporating known SNPs and short indels (Sanger REL-1505) into the reference mouse genome sequence (Genome Reference Consortium Mouse Reference 38) using `g2gtools` (http://churchill-lab.github.io/g2gtools/). We lifted the reference gene annotation (Ensembl Release 78) over to the CAST genome coordinates, and derived a CAST-specific transcriptome. The B6 transcriptome is based on the mouse reference genome. We constructed a bowtie (v1.0.0) index to represent the diploid transcriptome with two alleles of each transcript. We aligned reads using bowtie with parameters ‘–all’, ‘–best’, and ‘–strata’, allowing for 3 mismatches (‘-v 3’). These settings enable us to find all of the best alignments for each read. For example, if there is a zero-mismatch alignment for a read, all alignments with zero mismatch will be accepted.

### Overview of the scBASE model

The `scBASE` algorithm is composed of three steps: *read counting*, *classification*, and *estimation* (Figure 1). The read counting step is applied first to resolve read mapping ambiguity due to multi-reads and to estimate expected read counts. The read counting step is not a requirement since the following steps are applicable to any allele-specific count estimates. The classification and estimation steps are executed iteratively to classify the allelic expression state and to estimate the allelic proportions for each gene in each cell using a hierarchical mixture model. We have implemented `scBASE` as a Monte Carlo Markov chain (MCMC) algorithm^30^, which randomly samples parameter values from their conditional posterior distributions. We have also implemented the classification and estimation steps as an Expectation-Maximization (EM) algorithm^31^ that converges to the maximum a posteriori parameter estimates (Supplemental Methods). MCMC is flexible, and the sampling distributions and priors are easy to change in the MCMC code. MCMC provides the full posterior distribution of allelic proportions and thus provides useful information about the uncertainty of estimated parameters. We also found that MCMC is more stable when fitting allelic proportion of monoallelic classes. The EM algorithm is much faster, but it provides only point estimation. We provide a brief description of the algorithm here and provide additional details in Supplemental Methods.

### Read counting

In order to count all of the available sequence reads for each gene and allele, we have to resolve read mapping ambiguity that occur when aligning reads to a diploid genome. Genomic multi-reads align with equal quality to more than one gene. Allelic multi-reads align with equal quality to both alleles of a gene. In `scBASE`, multi-reads are resolved by computing a weighted allocation based on the estimated probability of each alignment. We use an EM algorithm implemented in `EMASE` software for this step^16^. Alternatively, read counting could be performed using similar methods implemented in `RSEM`^14^ or `kallisto`^15^ software. The estimated maternal read count (*x*_*gk*_) for each gene (*g*) in each cell (*k*) is the weighted sum of all reads that align to the maternal allele, where the weights are proportional to the probability of the read alignment. Similarly, the estimated paternal read count (*y*_*gk*_) is the weighted sum of all reads that align to the paternal allele. The total read count is the sum of the allele-specific counts (*n*_*gk*_ = *x*_*gk*_ + *y*_*gk*_). A parameter of interest is the allelic proportion *p*_*gk*_. The read counting step provides an initial estimate 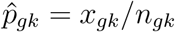, which we refer to as the weighted allocation estimated counts (ii).

### Classification

In the classification step, we estimate the allelic expression state (*z*_*gk*_) for each gene in each cell. The allelic expression state is a latent variable with three possible values *z_gk_ ∈ {P, B, M}* representing paternal monoallelic, bi-allelic, and maternal monoallelic expression, respectively. Uncertainty about the allelic expression state derives from sampling variation that can produce zero counts for one or both alleles even when the allele-specific transcripts may be present in the cell. We account for this uncertainty by computing a probabilistic classification based on a mixture model in which the maternal read counts *x*_*gk*_ are drawn from one of three beta-binomial distributions (given *n*_*gk*_) according to the allelic expression state *z*_*gk*_. For a gene in the bi-allelic expression state the maternal allelic proportion is denoted 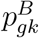 and, as suggested by the notation, it may vary from cell to cell following a beta distribution. For a gene in the paternal monoallelic expression state, the allelic proportion 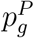 follows a beta distribution with a high concentration of mass near zero. Similarly, for a gene in the maternal monoallelic expression state, we model 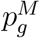 using a beta distribution with the concentration of mass near one. The beta distribution parameters for the maternal and paternal states are gene-specific but are constant across cells.

### Estimation

The classification step assumes that the mixture model parameters are known. This model describes gene-specific allelic proportions for each cell and thus it has a very large number of parameters. In the scRNA-Seq setting where thousands of genes are measured but low read counts and sampling zeros are prevalent, we may have limited data to support their reliable estimation. Bayesian analysis of the hierarchical model treats parameters as random variables and is well suited for this type of estimation. In this context, the hierarchical model improves the precision of estimation by borrowing information across cells for each gene, giving more weight to cells that are in the same allelic expression state. This estimation technique is referred to as *partial pooling*. Specifically, we sample the mixture weights 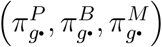 and the class-specific allele proportions 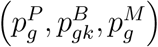; generate classification probabilities 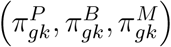; and then estimate the allelic proportions as a weighted average

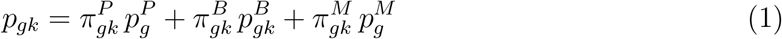

The average value across many iterations is 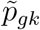, the partial pooling estimator.

### Estimating allelic proportions in subpopulations of cells or genes

The `scBASE` algorithm is designed to model heterogeneous ASE states in any population of cells. In some cases, as in the developmental series of Deng et al., it is of interest to focus on different subpopulations. When subpopulations of cells or groups of genes, e.g., X chromosome genes, are expected to have different distributions of allelic states, we recommend two options. The first option is to run the MCMC implementation of `scBASE` separately for each group. The strength of this approach is that it provides the posterior distribution of group-specific allelic proportions. However the level of uncertainty could increase for estimated parameters when the number of cells in any group is limited. The second option is to first run MCMC with all the available cells and estimate the prior parameters, 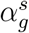 and 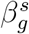. These prior parameters describe how allelic proportions are distributed in the monoallelic and bi-allelic states, and therefore, are common across all groups. Then using estimated values for these parameters, re-estimate the remaining parameters, 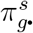, 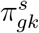, and 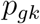, within each cell type using the EM algorithm. In the restricted version of EM, we iteratively update 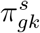 (E-step) and 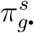 (M-step) for cells within each subpopulation. Once 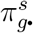 and 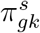 converge, we can compute 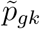 using Equation (1). We applied this second approach to Deng et al. time series data along mouse embryo development (n=286 cells). Genes on the X chromosome present another example where it makes sense to run `scBASE` separately, in this case on two subpopulations of genes. Our analyses of female X chromosome genes used this strategy (Supplemental Figures S9, S10, and S11).

### Assigning allelic expression states from estimated counts

Unique read counts are obtained directly from counting reads after discarding all genomic and allelic multi-reads. Weighted allocation counts are derived from the EM algorithm as described above. To estimate counts after partial pooling, we multiply 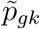 by the total gene expression counts. We note that estimated counts are not integers and may be non-zero but less than one. Classification of allelic expression states for each gene in each cell directly from observed or estimated counts requires setting a threshold for monoallelic expression. For each allele, we regarded it as expressed if its estimated abundance is greater than one reads (or one UMI as in Larsson et al^21^).

### Classification of a gene according to its ASE profile across many cells

We classify a gene according to the proportion of cells in P-, B-, and M-states, 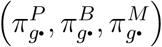, that are estimated by the partial pooling model. If a majority of cells 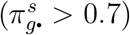 are in a particular ASE state, *s ∈ {P, B, M}*, then we will assign the gene to the class **P** (monoallelic paternal; blue), **B** (bi-allelic; yellow), or **M** (monoallelic maternal; red) respectively. When a majority of cells are a mixture of two of those classes (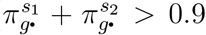 where *s*_1_*, s*_2_ *∈ {P, B, M}*), we classify it into either of **PB** (mixture of monoallelic paternal and bi-allelic; green), **BM** (mixture of monoallelic maternal and bi-allelic; orange), or **MP** (a mixture of monoallelic maternal and paternal; purple). Otherwise, genes that present all three ASE states are classified as **PBM** (mixture of all; gray). We specified these seven classes in a ternary simplex diagram (Figure 5a)^32^. The class boundaries are arbitrary but the aim of this classification is to provide a simple descriptive summary of the gene expression states present in a population of cells.

### Sampling reads

We randomly sampled 1% of reads in each of 122 cells at the early, mid, and late blastocyst stages to obtain an average read count of *∼*148k reads per cell. We chose the blastocyst cell types because, unlike cells in earlier developmental stages, they show the widest range of different states of allelic expression. The original analysis of `SCALE`^23^ also used the same 122 cells. We applied the unique-reads method and weighted allocation algorithm to the full set of *∼*14.8M reads and also applied each of four estimation methods (unique reads, weighted allocation counts, unique reads with partial pooling, and weighted allocation with partial pooling) to the down-sampled data. We compared estimates obtained from the down-sampled data to the full data estimates and computed the mean squared error of estimation across cells for each gene.

### Simulation of counts under perfect independence model

We randomly sampled the marginal probabilities of maternal and paternal allelic expression, *p*_*M*_ and *p*_*P*_ from uniform distribution between 0 and 1. Then we generated 2*×*2 tables by sampling counts from multinomial distribution with probability {*p*_*M*_*p*_*P*_, *p*_*M*_(1−*p*_*P*_), (1−*p*_*M*_)*p*_*P*_, (1−*p*_*M*_)(1−*p*_*P*_)} for bi-allelic, maternal monoallelic, paternal monoallelic, and silent cells respectively.

## Acknowledgments

We would like to thank Steven C. Munger and Daniel A. Skelly for their helpful comments on this manuscript.

## Author Contributions

KC, NR, and GC conceived and planned the study. KC performed the model implementation and analyses. KC and GC interpreted the scientific findings. KC and GC discussed, and wrote the manuscript.

