## Supplemental for "scBASE: A Bayesian mixture model for the analysis of allelic expression in single cells"

### Supplemental Methods

**MCMC algorithm.** We consider a single gene $\mathcal{G}$ in cells indexed by $k = 1, \ldots, K$. The maternal and paternal read counts, $x_{gk}$ and $y_{gk}$, are estimated by the weighted allocation algorithm or unique-reads method. The expected counts are non-integral in many cases, we convert them to integer with floor function. The total read count for $\mathcal{G}$ is $n_{gk} \equiv x_{gk} + y_{gk}$. We define the maternal allelic proportion $p_{gk}$ for each cell to be the expected proportion of maternal reads. We assume each cell is in one of three states with respect to the expression of $\mathcal{G}$: $s \in \{P, B, M\}$ where the indices denote $(P)$aternal monoallelic, $(B)$i-allelic, or $(M)$aternal monoallelic expression respectively. For each cell $k$, we introduce an indicator vector, $\boldsymbol{z}_{gk} = \left(z_{gk}^{P},\, z_{gk}^{B},\, z_{gk}^{M}\right)$ where

$$
\begin{aligned}
z_{gk}^{s} &= \begin{cases} 1, & \text{if } \mathcal{G} \text{ is in expression state } s, \\ 0, & \text{otherwise} \end{cases} \\
\text{and } \sum_{s} z_{gk}^{s} &= 1.
\end{aligned}
\tag{1}
$$

By definition, the marginal distribution of $P(z_{gk}^{s} = 1) = \pi_{g\bullet}^{s}$ and $\sum_{s} \pi_{g\bullet}^{s} = 1$ and we interpret the mixture weights, $\pi_{g\bullet}^{s}$, to be the proportion of cells in allelic expression state $s$ in the population. We propose a hierarchical mixture model:

$$
\begin{aligned}
x_{gk} &\sim \begin{cases} \text{Binomial}(n_{gk}, p_{g}^{P}) & \text{if } z_{gk}^{P} = 1, \\ \text{Binomial}(n_{gk}, p_{gk}^{B}) & \text{if } z_{gk}^{B} = 1, \\ \text{Binomial}(n_{gk}, p_{g}^{M}) & \text{if } z_{gk}^{M} = 1 \end{cases} \\
p_{g}^{P} &\sim \text{Beta}(1, \alpha_{g}^{mono}) \\
p_{gk}^{B} &\sim \text{Beta}(\alpha_{g}^{B}, \beta_{g}^{B}) \\
p_{g}^{M} &\sim \text{Beta}(\alpha_{g}^{mono}, 1) \\
\alpha_{g}^{mono} &\sim \text{half-Cauchy}(7, 2) \\
\alpha_{g}^{B},\, \beta_{g}^{B} &\sim \text{half-Cauchy}(2, 2) \\
\pi_{g\bullet}^{s} &\sim \text{Dirichlet}\left(\frac{1}{3}, \frac{1}{3}, \frac{1}{3}\right)
\end{aligned}
\tag{2}
$$

As analytical solution to the maximum likelihood estimators does not exist for these models, we implemented Markov-Chain Monte Carlo algorithm using `STAN`[30] version 2.17.0. While fitting `scBASE` model, we calculate $\pi_{gk}^{s}$

$$\pi_{gk}^{P} = E(z_{gk}^{P}=1) = \frac{\pi_{g\bullet}^{P}\,\text{Binomial}(x_{gk}|n_{gk}, p_g^P)}{N}$$

$$\pi_{gk}^{B} = E(z_{gk}^{B}=1) = \frac{\pi_{g\bullet}^{B}\,\text{Binomial}(x_{gk}|n_{gk}, p_{gk}^B)}{N} \quad \text{and}$$

$$\pi_{gk}^{M} = E(z_{gk}^{M}=1) = \frac{\pi_{g\bullet}^{M}\,\text{Binomial}(x_{gk}|n_{gk}, p_g^M)}{N} \quad \text{where} \tag{3}$$

$$\begin{aligned} N = {} & \pi_{g\bullet}^{P}\,\text{Binomial}(x_{gk}|n_{gk}, p_g^P) + \\ & \pi_{g\bullet}^{B}\,\text{Binomial}(x_{gk}|n_{gk}, p_{gk}^B) + \\ & \pi_{g\bullet}^{M}\,\text{Binomial}(x_{gk}|n_{gk}, p_g^M) \end{aligned}$$

We then calculated $p_{gk}$ in the following way.

$$p_{gk} = \pi_{gk}^{P}\,p_g^P + \pi_{gk}^{B}\,p_{gk}^B + \pi_{gk}^{M}\,p_g^M \tag{4}$$

Once model fitting finishes, we report a maximum a posteriori estimate of maternal allele proportion, $\tilde{p}_{gk}$, as the mean of sampled $p_{gk}$. We provide our `STAN` implementation below.

**The three-component Beta-Binomial mixture model.**

---

```
data {
   int<lower=1> N;
   int<lower=0> n[N];
   int<lower=0> x[N];
}
parameters {
   simplex[3] pi;
   real<lower=7> a_mono;
   vector<lower=2>[2] alpha;
   vector<lower=0,upper=1>[N] theta;
}
model {
   vector[3] log_pi = log(pi);
   a_mono ~ cauchy(7, 2);
   alpha ~ cauchy(2, 2);
   theta ~ beta(alpha[1], alpha[2]);
   for (i in 1:N) {
      vector[3] lps = log_pi;
      lps[1] = lps[1] + beta_binomial_lpmf(x[i]|n[i], a_mono, 1);
```

```
      lps[2] = lps[2] + beta_binomial_lpmf(x[i]|n[i], 1, a_mono);
      lps[3] = lps[3] + binomial_lpmf(x[i]|n[i], theta[i]);
      target += log_sum_exp(lps);
   }
}
generated quantities {
   matrix[N,3] pi_z;
   real log_sum_exp_log_pi_z_raw;
   for (i in 1:N) {
      vector[3] log_pi_z_raw = log(pi);
      log_pi_z_raw[1] += beta_binomial_lpmf(x[i]|n[i], a_mono, 1);
      log_pi_z_raw[2] += beta_binomial_lpmf(x[i]|n[i], 1, a_mono);
      log_pi_z_raw[3] += binomial_lpmf(x[i]|n[i], theta[i]);
      log_sum_exp_log_pi_z_raw = log_sum_exp(log_pi_z_raw);
      for (j in 1:3)
         pi_z[i,j] = exp(log_pi_z_raw[j] - log_sum_exp_log_pi_z_raw);
   }
}
```

---

**Expectation-Maximization algorithm.** For point estimation of maternal allele proportion in each cell, $p_{gk}$, for gene $\mathcal{G}$, we also propose an empirical Bayes model based on the same latent variable, $\boldsymbol{z}_{gk} = (z_{gk}^P, z_{gk}^B, z_{gk}^M)$:

$$\begin{aligned} x_{gk}\Big|p_{gk}, n_{gk} &\sim \text{Binomial}(n_{gk}, p_{gk}) \\ p_{gk}\Big|z_{gk}^s &\sim \text{Beta}(\alpha_g^s, \beta_g^s) \\ \boldsymbol{z}_{gk} &\sim \text{Multinomial}\Big(1, \{\pi_{g\bullet}^P, \pi_{g\bullet}^B, \pi_{g\bullet}^M\}\Big) \end{aligned} \tag{5}$$

We note that $\alpha_g^s$ and $\beta_g^s$ can be specified and fixed in which case the following model reduces to a simpler form that does not update them along the EM process. The distribution of $x_{gk}\Big|n_{gk}, z_{gk}^s$ is Beta-Binomial and the full marginal distribution of $x_{gk}\Big|n_{gk}$ is 3-component Beta-Binomial mixture:

$$\begin{aligned} P(x_{gk}\Big|n_{gk}) &= \sum_s P(z_{gk}^s = 1)\, P(x_{gk}|n_{gk}, z_{gk}^s{=}1) \\ &= \sum_s \pi_{g\bullet}^s \,\text{Beta-Binomial}\Big(x_{gk}\Big|n_{gk}, \alpha_g^s, \beta_g^s\Big). \end{aligned} \tag{6}$$

The log likelihood is

$$
\begin{aligned}
\log L(\boldsymbol{X}_g|\boldsymbol{N}_g) &= \log P(x_{g1},\ldots,x_{gK}\Big|n_{g1},\ldots,n_{gK}) \\
&= \log \prod_k P(x_{gk}\Big|n_{gk}) \\
&= \sum_k \log \sum_s \pi^s_{g\bullet} P(x_{gk}|n_{gk}, z^s_{gk}=1) \\
&= \sum_k \log \sum_s \pi^s_{g\bullet} \binom{n_{gk}}{x_{gk}} \frac{B\Big(\alpha^s_g + x_{gk},\ \beta^s_g + n_{gk} - x_{gk}\Big)}{B\Big(\alpha^s_g, \beta^s_g\Big)} \\
\text{where } B(\alpha,\beta) &= \frac{\Gamma(\alpha)\Gamma(\beta)}{\Gamma(\alpha+\beta)}.
\end{aligned}
\tag{7}
$$

As analytical solution to the maximum likelihood estimators does not exist, we use the following Expectation-Maximization (EM) algorithm.

**E-step:** Given parameters $\alpha^s_g$, $\beta^s_g$, $\pi^s_{g\bullet}$, and data $x_{gk}$ and $n_{gk}$, we can compute, $\pi^s_{gk}$, the posterior expectation on $z^s_{gk}$.

$$
\begin{aligned}
\pi^s_{gk} &= E(z^s_{gk}=1|x_{gk}, n_{gk}, \widehat{\boldsymbol{\alpha}}_g, \widehat{\boldsymbol{\beta}}_g, \widehat{\boldsymbol{\pi}}_g) \\
&= \frac{P(z^s_{gk}=1)P(x_{gk}|n_{gk}, z^s_{gk}{=}1)}{\sum_{t\in\{P,B,M\}} P(z^t_{gk}=1)P(x_{gk}|n_{gk}, z^t_{gk}{=}1)} \\
&= \frac{\widehat{\pi}^s_{g\bullet} P(x_{gk}|n_{gk}, z^s_{gk}{=}1)}{\sum_t \widehat{\pi}^t_{g\bullet} P(x_{gk}|n_{gk}, z^t_{gk}{=}1)} \\
\text{where } \ \widehat{\boldsymbol{\alpha}}_g &= \left\{\widehat{\alpha}^P_g, \widehat{\alpha}^B_g, \widehat{\alpha}^M_g\right\}, \widehat{\boldsymbol{\beta}}_g = \left\{\widehat{\beta}^P_g, \widehat{\beta}^B_g, \widehat{\beta}^M_g\right\}, \widehat{\boldsymbol{\pi}}_g = \left\{\widehat{\pi}^P_{g\bullet}, \widehat{\pi}^B_{g\bullet}, \widehat{\pi}^M_{g\bullet}\right\}, \text{ and,} \\
P\Big(x_{gk}\Big|n_{gk}, z^s_{gk}{=}1\Big) &= \binom{n_{gk}}{x_{gk}} \frac{B\Big(\alpha^s_g + x_{gk},\ \beta^s_g + n_{gk} - x_{gk}\Big)}{B\Big(\alpha^s_g, \beta^s_g\Big)}.
\end{aligned}
\tag{8}
$$

**M-step:** Given the expectation on expression state of gene $\mathcal{G}$ in each cell, $\pi^s_{gk}$, we can compute the maximum likelihood estimators, $\widehat{\alpha}^s_g$, $\widehat{\beta}^s_g$, and $\widehat{\pi}^s_g$. What makes the maximum likelihood process more complicated in the proposed model is that $n_{gk}$'s are different cell by cell, and we want to put more weights on cells that have larger $n_{gk}$'s. Since $\pi^s_{gk}$ describes how likely cell $k$ belong to expression state $s$, we can make cells that are more likely to be in state $s$ to contribute more in estimating parameters $\alpha^s_g$, $\beta^s_g$, and $\pi^s_g$. We used the method of moments approach[31] in which Kleinman et al. equates the weighted mean and variance of observed maternal allele proportion, $x_{gk}/n_{gk}$, to the analytical mean and variance of Beta-Binomial distribution re-parameterized with $\mu_g^s = \alpha_g^s/(\alpha_g^s + \beta_g^s)$ and $\tau_g^s = (1 + \alpha_g^s + \beta_g^s)^{-1}$:

$$\begin{aligned} E\left(\frac{x_{gk}}{n_{gk}}\middle| z_{gk}^s{=}1\right) &= \mu_g^s \\ Var\left(\frac{x_{gk}}{n_{gk}}\middle| z_{gk}^s{=}1\right) &= \frac{\mu_g^s(1-\mu_g^s)}{n_{gk}} + \tau_g^s \mu_g^s (1-\mu_g^s)\left(1 - \frac{1}{n_{gk}}\right) \end{aligned} \tag{9}$$

Then, parameters are updated in the following way.

$$\begin{aligned} \left(\widehat{\mu}_g^s\right)^{new} &= \bar{p}_g^s \\ \left(\widehat{\tau}_g^s\right)^{new} &= \frac{S_g^s - \bar{p}_g^s(1-\bar{p}_g^s)\left\{\sum_k \frac{w_{gk}^s}{n_{gk}}\left(1 - \frac{w_{gk}^s}{W_g^s}\right)\right\}}{\bar{p}_g^s(1-\bar{p}_g^s)\left\{\sum_k w_{gk}^s\left(1 - \frac{w_{gk}^s}{W_g^s}\right) - \sum_k \frac{w_{gk}^s}{n_{gk}}\left(1 - \frac{w_{gk}^s}{W_{gk}^s}\right)\right\}} \end{aligned} \tag{10}$$

where

$$\begin{aligned} \bar{p}_g^s &= \frac{\sum_k w_{gk}^s \left(\frac{x_{gk}}{n_{gk}}\right)}{W_g^s} \\ S_g^s &= \sum_k w_{gk}^s \left(\left(\frac{x_{gk}}{n_{gk}}\right) - \bar{p}_g^s\right)^2, \\ w_{gk}^s &= \pi_{gk}^s \cdot \frac{n_{gk}}{1 + (n_{gk}-1)\left(\widehat{\tau}_g^s\right)^{old}}\ , \quad \text{and} \\ W_g^s &= \sum_k w_{gk}^s \end{aligned} \tag{11}$$

From updated $\mu_g^s$ and $\tau_g^s$, we can easily derive the parameters that are used in (5).

$$
\begin{aligned}
\left(\widehat{\alpha}_g^s\right)^{new} &= \left\{ \frac{1}{\left(\widehat{\tau}_g^s\right)^{new}} - 1 \right\} \left(\widehat{\mu}_g^s\right)^{new}, \text{ and} \\
\left(\widehat{\beta}_g^s\right)^{new} &= \left( \frac{1}{\left(\widehat{\tau}_g^s\right)^{new}} - 1 \right) \left( 1 - \left(\widehat{\mu}_g^s\right)^{new} \right)
\end{aligned} \tag{12}
$$

Here we designed the weight of each cell, $w_{gk}^s$, to reflect both $\pi_{gk}^s$ and $1/\tau_g^s$. By definition, $\pi_{gk}^s$ gets larger when the gene $\mathcal{G}$ in $k$-th cell is more likely to be in expression state $s$. Similarly, $1/\tau_g^s$ increases if the observed allele proportion, $x_{gk}/n_{gk}$, reside in narrower range across the cell population, which positively correlates to the sample sizes, $n_{gk}$. Therefore, cells that are more likely to be in state $s$ and also having higher depth of coverage would contribute more on estimating parameter $\alpha_g^s$ and $\beta_g^s$. Finally, we update $\pi_{g\bullet}^s$ by adding the expected expression state of $\mathcal{G}$ across the cell population.

$$
\left(\widehat{\pi}_{g\bullet}^s\right)^{new} = \frac{1}{K} \sum_k \pi_{gk}^s \tag{13}
$$

After parameter values converge, we also report the posterior expected allele proportion, $\tilde{p}_{gk}(\equiv \tilde{x}_{gk}/n_{gk})$. We show here that the posterior allele proportion of cell $k$ lies in-between what is observed in cell $k$ itself and what is found from other cells in the same expression state $s$.

$$
\begin{aligned}
\tilde{p}_{gk} &= E\Big(x_{gk} \Big| n_{gk}\Big) \\
&= E\Big(\sum_s \pi_{g\bullet}^s \,\text{Beta-Binomial}\Big(x_{gk} \Big| n_{gk}, \alpha_g^s, \beta_g^s\Big)\Big) \\
&= \sum_s \pi_{g\bullet}^s \frac{\widehat{\alpha}_g^s}{\widehat{\alpha}_g^s + \widehat{\beta}_g^s}
\end{aligned} \tag{14}
$$

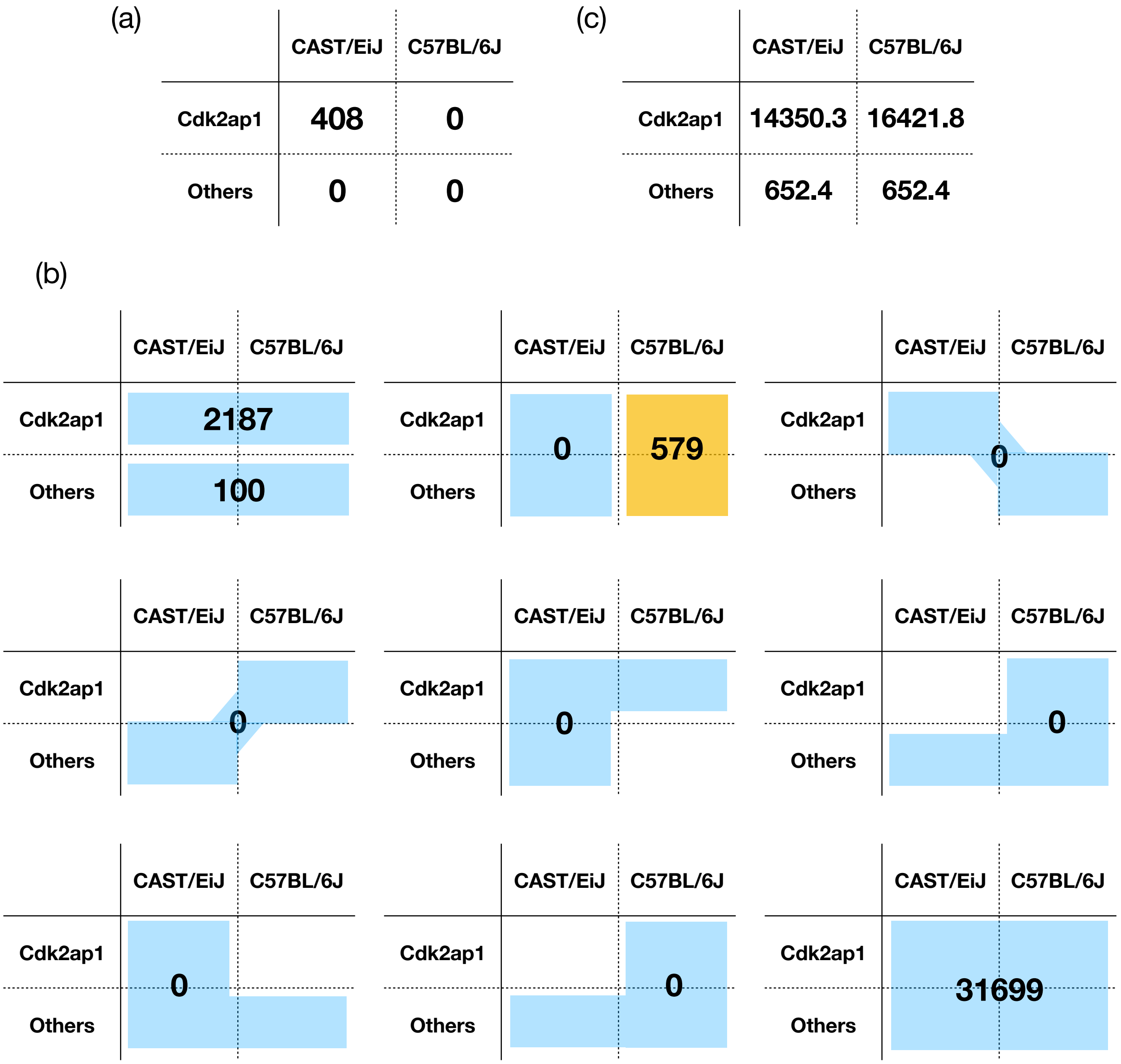


Supplemental Figure S1: Allele-specific multi-reads. We examined the read count data for *Cdk2ap1* in a single cell from the 16-cell stage of development. **(a)** There are 408 unique reads and all of them align to the CAST (maternal) allele of *Cdk2ap1*, which is consistent with monoallelic expression of this gene. **(b)** We enumerated the reads that fall into each of the possible multi-alignment patterns. Among the allelic multi-reads, there are 2,187 reads that align equally well to both alleles of *Cdk2ap1* and 100 reads aligned uniquely to both alleles of another gene family member. There are 579 genomic multi-reads and all of these align to the B6 allele of *Cdk2ap1* and at least one other gene. The remaining 31,699 reads align equally well to both alleles of *Cdk2ap1* and at least one other gene family member. The allelic and genomic multi-reads provide information needed to estimate the expected distribution of alignments for all of the reads. **(c)** The weighted allocation algorithm predicts that *Cdk2ap1* in expressed as a bi-allelic gene with a maternal allelic proportion $\hat{p}_{gk} = 0.466$.

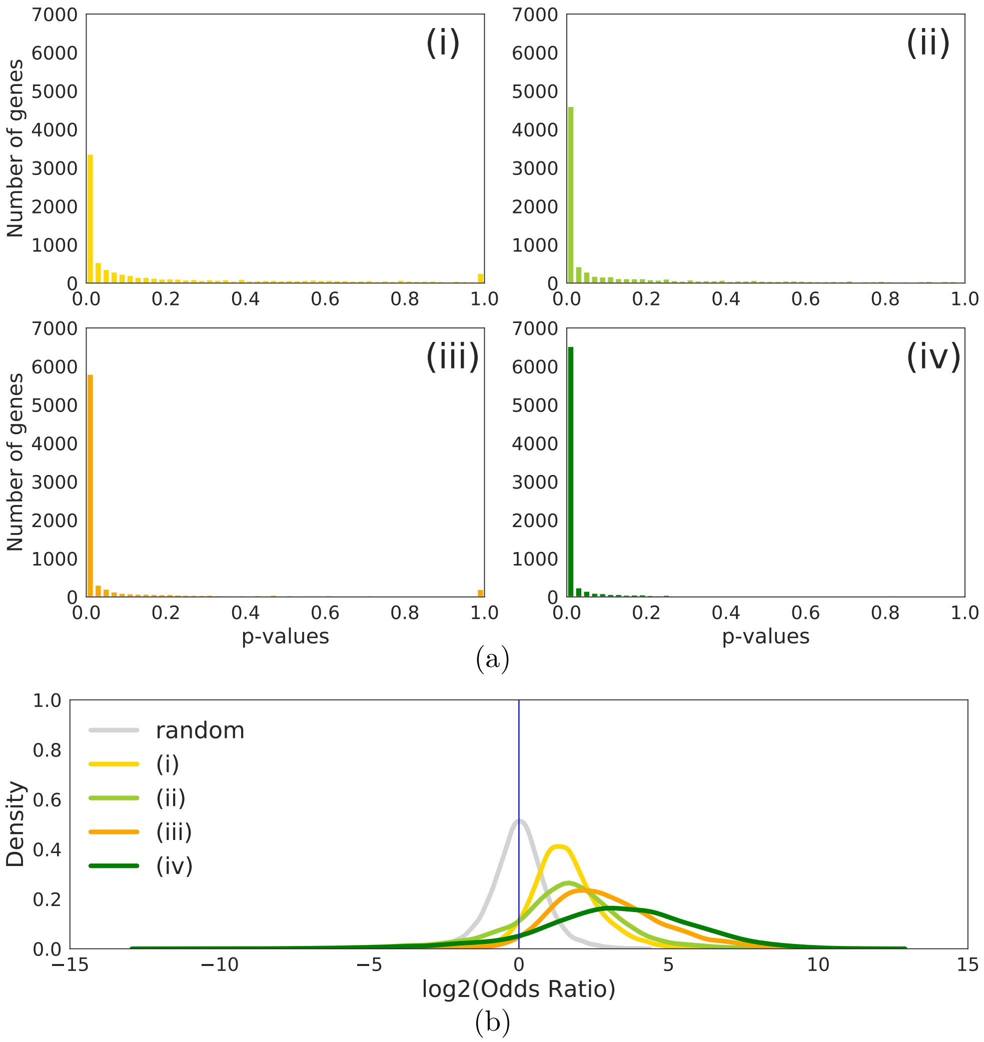


Supplemental Figure S2: **(a)** The distribution of p-values for testing independence of allelic bursting with 122 mature blastocyst cells[7]. We evaluated the significance of the independence test using the log-odds ratio (logOR) as implemented in the `SCALE` software[23] (`non_ind_bursting` function) using (i) unique reads, (ii) weighted allocation, (iii) unique reads with partial pooling, and (iv) weighted allocation with partial pooling. In all cases, a majority of genes show strong evidence for non-independent bursting. `SCALE` reports 3,381 genes that are non-independent using the results of unique-reads method, 4,815 genes using weighted allocation, 6,068 genes by unique reads with partial pooling, and 6,761 genes based on weighted allocation with partial pooling at the FDR level of 5%. **(b)** In addition, thousands of genes had $|\text{logOR}|>2$, for example, 2,845 and 3,763 out of 8,290 genes using unique reads and weighted allocation. More genes have $|\text{logOR}|>2$ after partial pooling with `scBASE`: 5,622 and 6,209 respectively. This result is also consistent with a strong tendency for allelic bursts to occur synchronously.

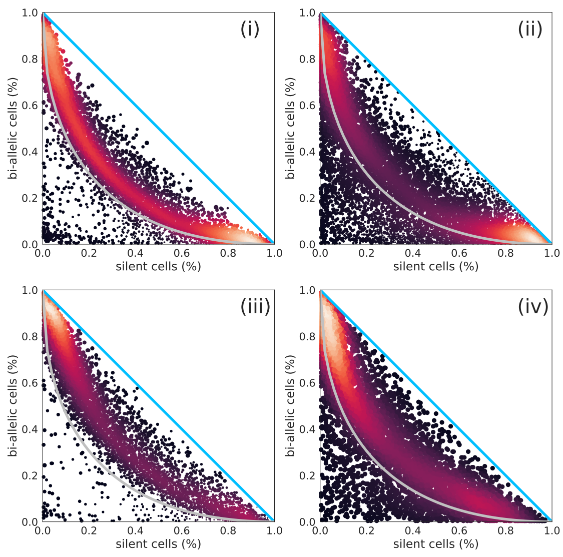


(a)

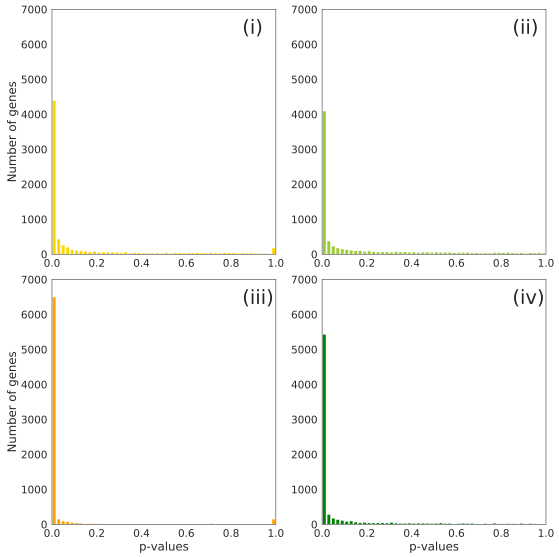


(b)

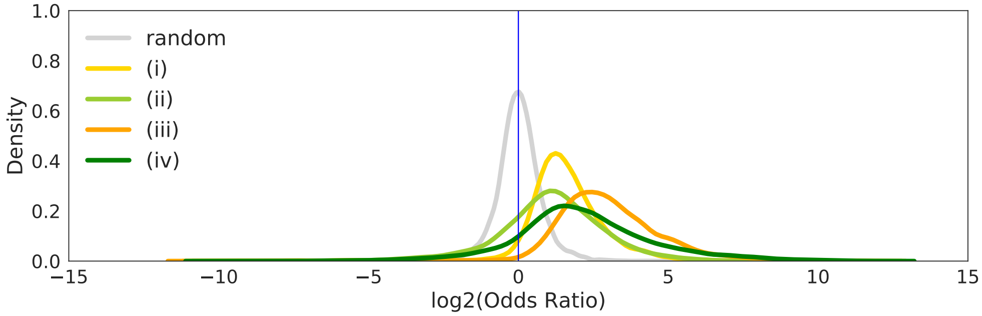


(c)

Supplemental Figure S3: **(a)** Tests for independence of allelic bursting with 238 mouse primary fibroblast cells[24]. We estimated proportions of bi-allelic and silent cells as proposed in Deng et al.[7] using counts based on (i) unique reads, (ii) weighted allocation, (iii) unique reads with partial pooling, and (iv) weighted allocation with partial pooling. **(b)** In contrast to simulated data based on the independence model in which most genes appear below the perfect independence curve (Figure 4b), most genes lie above the curve. We conclude that Deng et al. approach, when correctly interpreted, supports non-independence of allelic bursting. A non-independence test implemented in `SCALE` (`non_ind_bursting` function) identified that (i) 4,643 genes are significantly non-independent based on unique reads, (ii) 4,262 genes by weighted allocation count, (iii) 6,677 genes by unique reads with partial pooling, and (iv) 5,682 genes based on weighted allocation with partial pooling at the FDR level of 5%. **(c)** This results agree with our conclusion based upon Deng et al. approach. In addition, most genes displayed significantly positive values of logOR[20, 22] irrespective of ASE quantification models. Thousands of genes had |logOR|>2, for example, 2,845 and 2,615 out of 7,757 genes using unique reads and weighted allocation. More genes have |logOR|>2 after partial pooling with `scBASE`: 5,736 and 4,143 respectively. This result is consistent with a strong tendency for allelic bursts to occur synchronously.

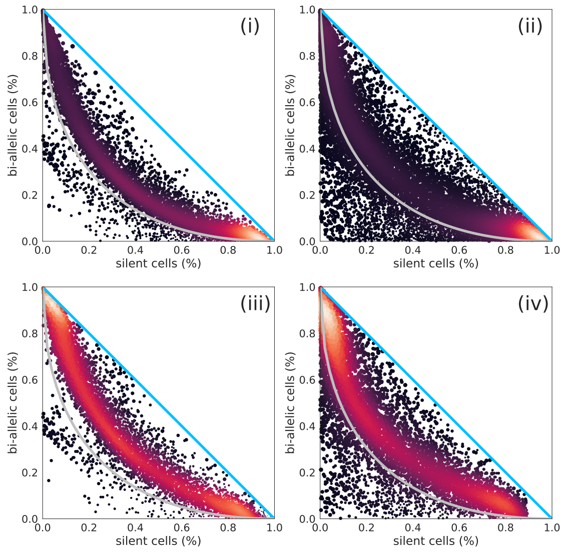


(a)

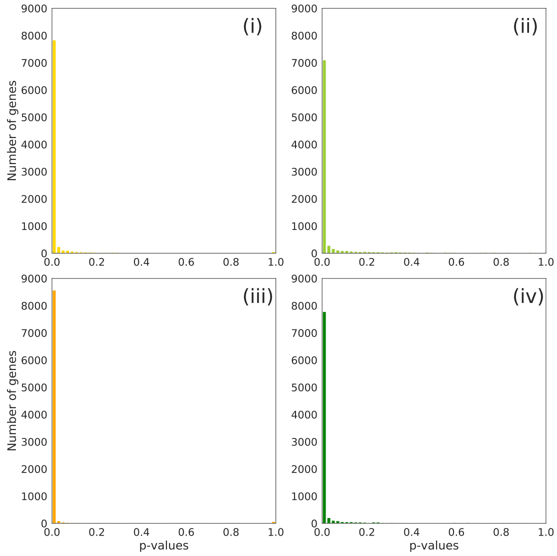


(b)

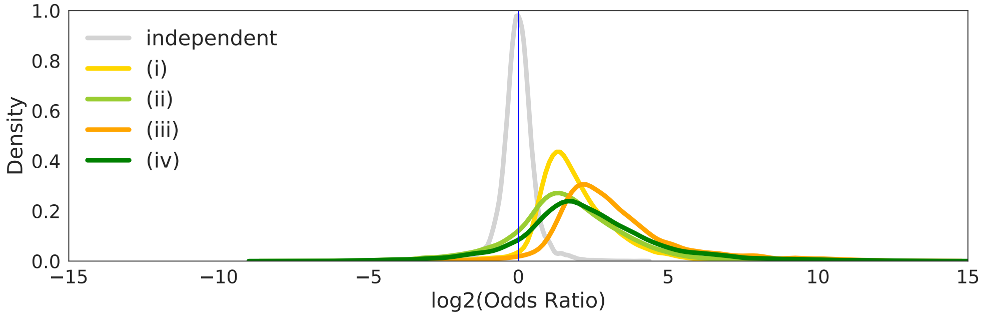


(c)

Supplemental Figure S4: **(a)** Tests for independence of allelic bursting with 534 mouse embryonic stem cells[25]. We estimated proportions of bi-allelic and silent cells as proposed in Deng et al.[7] using counts based on (i) unique reads, (ii) weighted allocation, (iii) unique reads with partial pooling, and (iv) weighted allocation with partial pooling. **(b)** In contrast to simulated data under the independence model in which most genes appear below the perfect independence curve (Figure 4b), most genes lie above the curve. We conclude that Deng et al. approach, when correctly interpreted, supports non-independence of allelic bursting. A non-independence test implemented in `SCALE` (`non_ind_bursting` function) identified that (i) 8,095 genes are significantly non-independent based on unique reads, (ii) 7,384 genes by weighted allocation, (iii) 8,682 genes by unique reads with partial pooling, and (iv) 8,008 genes based on weighted allocation with partial pooling at FDR level of 5%. **(c)** This results agree with our conclusion based upon Deng et al. approach. In addition, most genes displayed significantly positive values of logOR[20, 22] irrespective of ASE quantification models. Thousands of genes had |logOR|>2, for example, 3,751 and 3,950 out of 9,320 genes using unique reads and weighted allocation. More genes have |logOR|>2 after partial pooling with `scBASE`: 6,884 and 5,050 respectively. This result is consistent with a strong tendency for allelic bursts to occur synchronously.

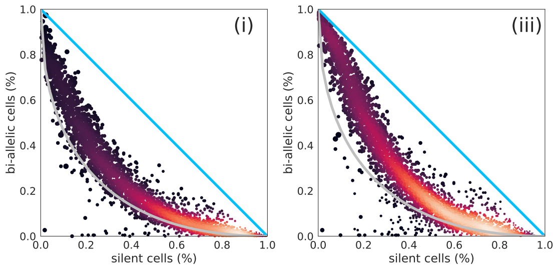


(a)

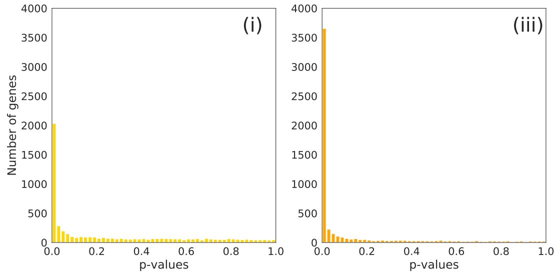


(b)

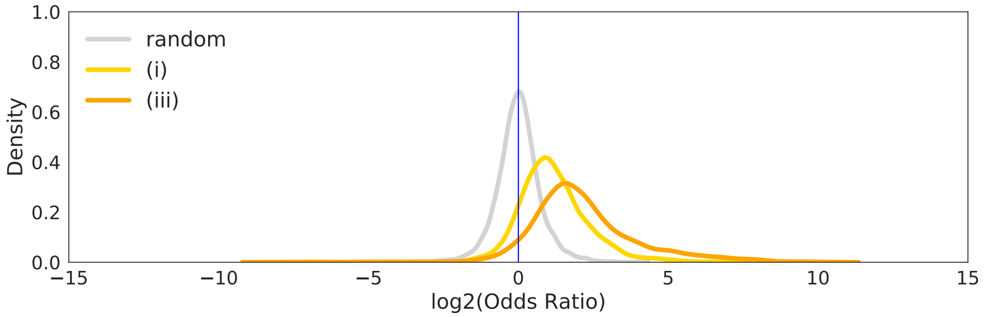


(c)

Supplemental Figure S5: **(a)** Tests for independence of allelic bursting with 224 primary mouse fibroblast cells[21]. Since this data set is generated from *custom* SMART-Seq2 platform, we downloaded the counts directly from https://github.com/sandberg-lab/txburst/blob/master/ExtDataFig1i.ipynb to perform the independence/non-independence tests. We estimated proportions of bi-allelic and silent cells using (i) original counts by Larsson et al.[21] and (iii) original counts with partial pooling. **(b)** Again most genes lie above the perfect independence curve, which does not support independence of allelic bursting. An independence test implemented in `SCALE` (`non_ind_bursting` function) identified that (i) 1,999 genes are significantly non-independent based on the original count, and (iii) 3,827 genes based on the original counts with partial pooling at the FDR level of 5%. **(c)** This results agree with our conclusion based upon Deng et al. approach. Most genes also displayed significantly positive values of logOR irrespective of ASE quantification models. Thousands of genes had |logOR|>2, for example, 1,011 out of 5,556 genes using the Larsson et al. original counts. 2,591 genes have |logOR|>2 after partial pooling with `scBASE`. This result is consistent with a strong tendency for allelic bursts to occur synchronously.

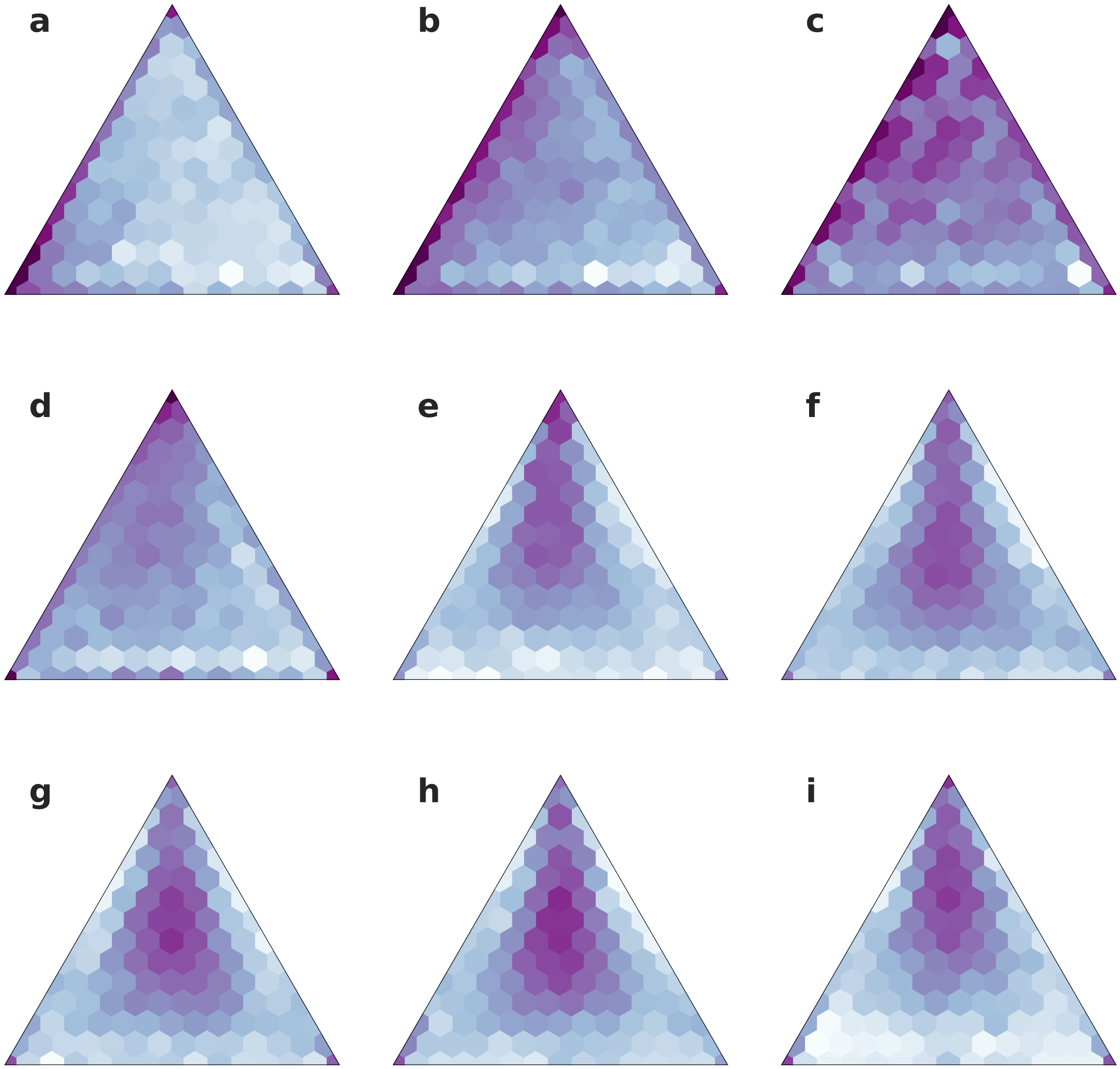


Supplemental Figure S6: Heatmaps show the distribution of genes over the simplex diagram for **a** the zygote & early 2-cell, **b** mid 2-cell, **c** late 2-cell, **d** 4-cell, **e** 8-cell, **f** 16-cell, **g** early blastocyst, **h** mid blastocyst, and **i** late blastocyst stage of development (See Figure 5b).

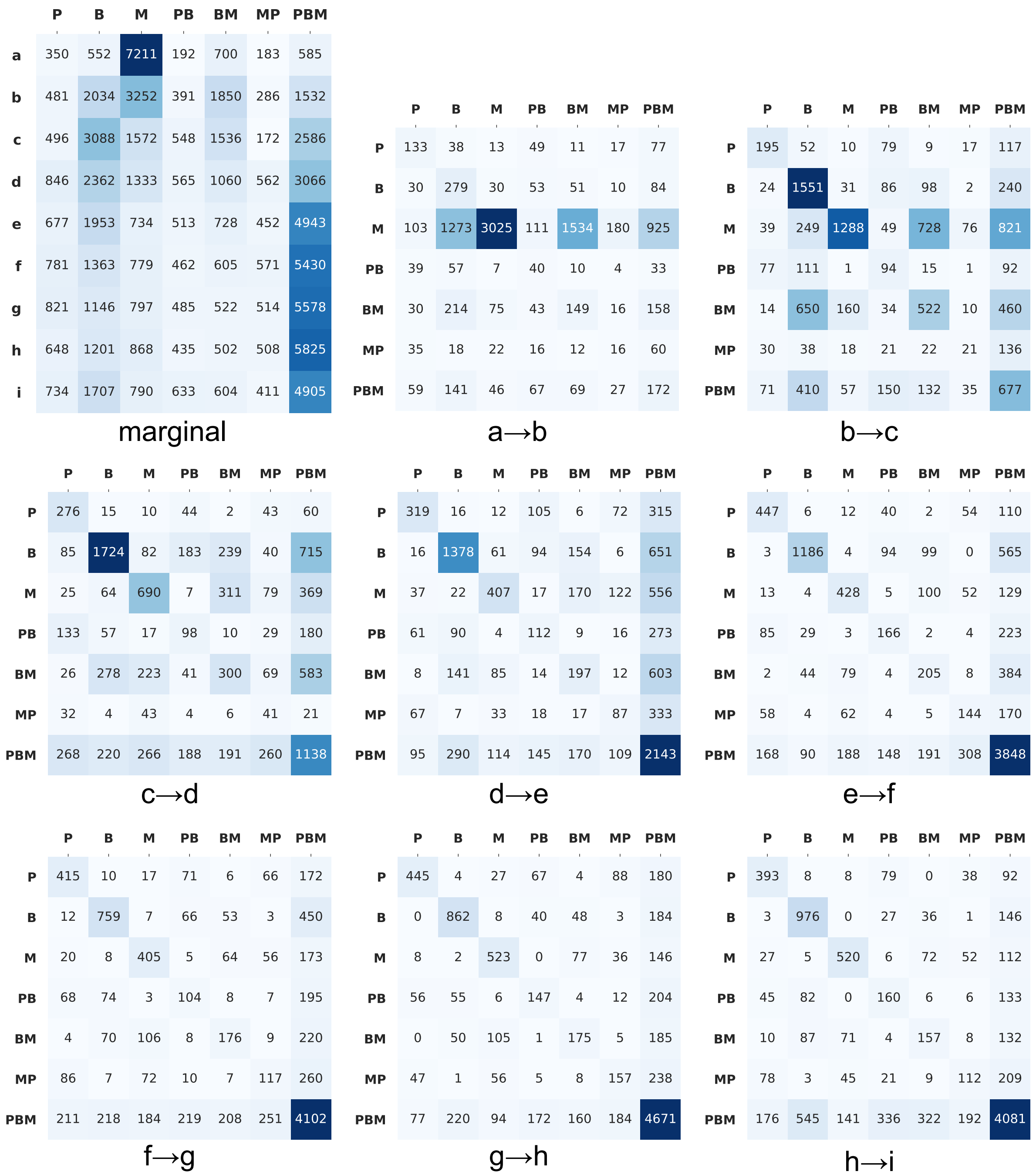


Supplemental Figure S7: The number of genes in each class and their transition among classes along the embryonic differentiation stages (See Figure 5b).

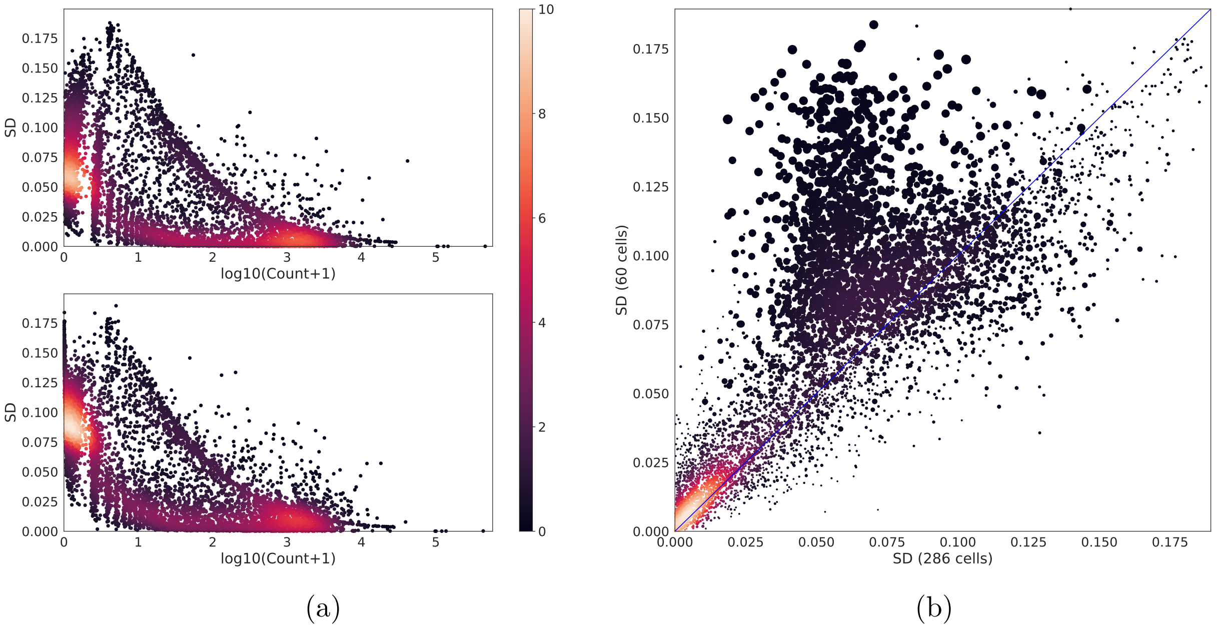


Supplemental Figure S8: Standard deviation (SD) of the estimated allelic proportions. We computed the SD of the partial pooling estimates of allelic proportions using all 286 cells and again using only 60 mid-blastocyst cells. We selected a representative mid-blastocyst cell, GSM1112706, to display the influence of using more cells on estimating allelic proportion of genes in each cell. **(a)** The SD based on pooling 286 cells (upper panel) and based on 60 cells (lower panel) is plotted against read coverage of ∼13,000 genes. We see the expected decrease in SD with increasing read coverage. The distinct upper and lower bands in these plots are due to the higher average SD for bi-allelic expressed genes versus monoallelic expressed genes. The SD is expected to be higher for bi-allelic genes because their allelic proportions are further from zero and one. **(b)** We directly compared the SDs obtained from 286 versus 60 cells. We observed generally higher SD for the 60 cell estimates and the effect is greatest when read coverage is low – point size in (b) is inversely proportional to log of read coverage. We conclude that increasing read depth and increasing cell number both improve the precision of the partial pooling estimates of allelic proportions.

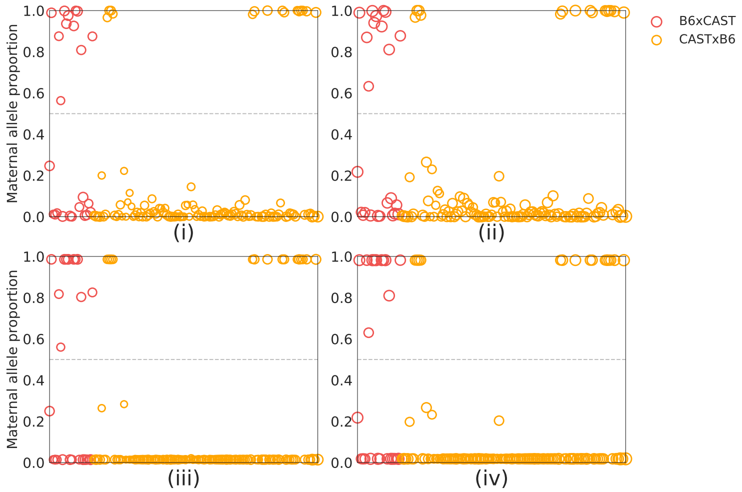


(a)

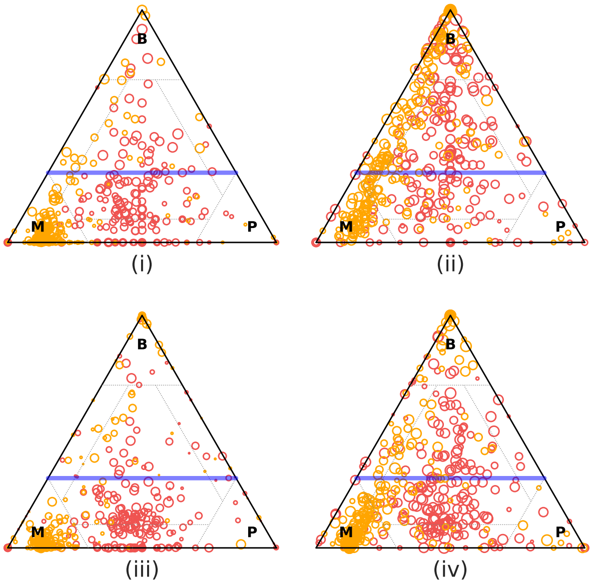


(b)

Supplemental Figure S9: X chromosome inactivation (XCI) in female primary fibroblast cells[24]. **(a)** We estimated maternal allele proportion of Xist gene across 145 female cells from the reciprocal cross of B6×CAST and CAST×B6 using (i) unique reads, (ii) weighted allocation, (iii) unique reads with partial pooling, and (iv) weighted allocation with partial pooling. Partial pooling corrects allelic proportion of Xist towards either of maternal or paternal monoallelic expression in many cells. **(b)** X chromosome genes are expressed from maternal allele in a large proportion of CAST×B6 cells for which Xist is expressed mostly from paternal allele (orange). On the other hand, Xist is randomly expressed from either of maternal and paternal alleles in B6×CAST cells, and X chromosome genes are enriched in **MP**-class (at the center below the blue line). ASE states of X chromosome genes are consistent to Xist allele-specific expression. Partial pooling also corrects the profiling of ASE states for both unique reads and weighted allocation, and strengthens the ASE pattern conforming to the XCI onset.

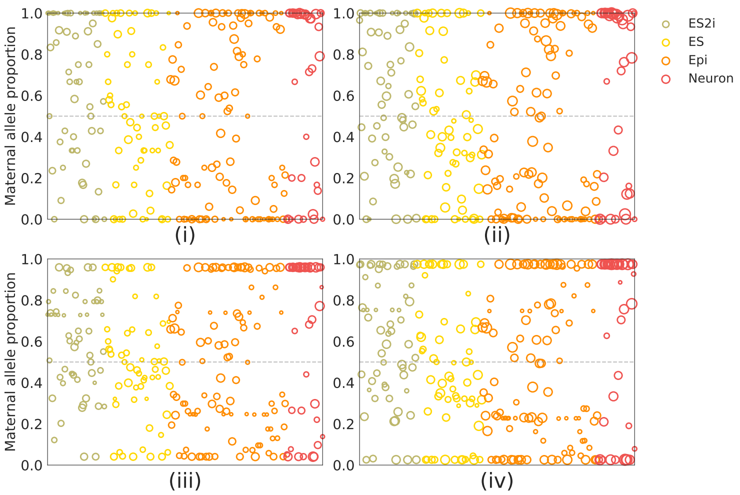


(a)

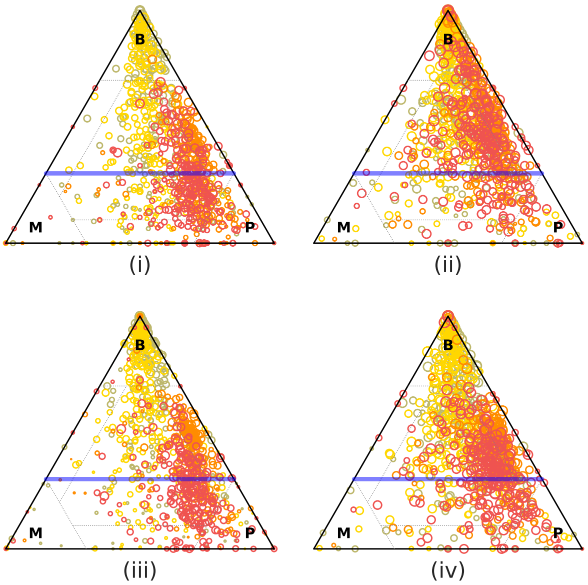


(b)

Supplemental Figure S10: X chromosome inactivation in female mESC, mEpiSC, and neuronal cells[25]. We estimated maternal allele proportion of Xist gene across 128, 120, and 39 female cells respectively using (i) unique reads, (ii) weighted allocation, (iii) unique reads with partial pooling, and (iv) weighted allocation with partial pooling. **(a)** Partial pooling corrects allele proportion of Xist towards either of monoallelic expression, but Xist is still expressed from both alleles in many cells. X chromosome genes are expressed from paternal allele in a high proportion of mEpiSCs and neuronal cells for which Xist is expressed largely from maternal allele (orange and red). **(b)** Allelic expression state of X chromosome genes are consistent to Xist allele expression. Partial pooling also corrects the Allelic expression state of these genes and displays paternal allele preference more clearly for both unique reads and weighted allocation, although overall incompleteness of XCI is common in mESCs, mEpiSCs, or neuronal cells irrespective of ASE estimation methods.

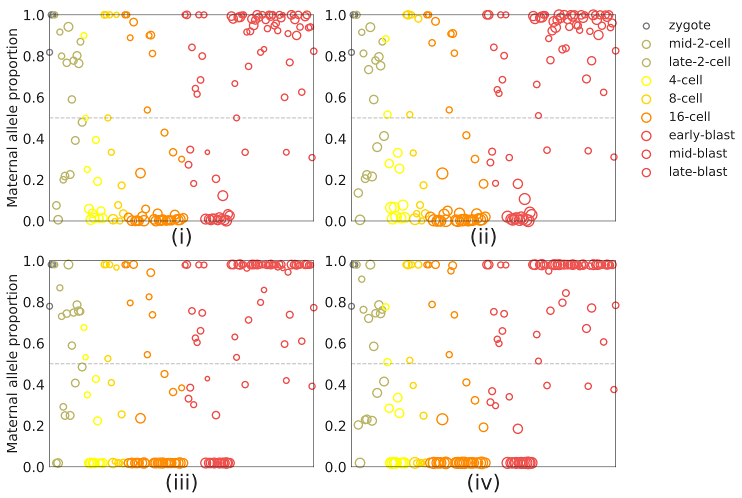


(a)

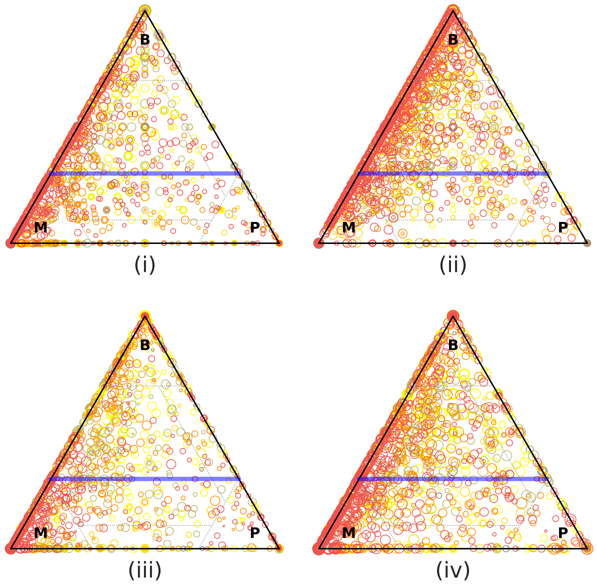


(b)

Supplemental Figure S11: X chromosome inactivation (XCI) in female pre-implantation blastocyst cells[7]. We sex-typed the cells based on Xist gene expression (>10 reads) and identified 155 female cells. We estimated maternal (CAST) allele proportion using (i) unique reads, (ii) weighted allocation, (iii) unique reads with partial pooling, and (iv) weighted allocation with partial pooling. **(a)** Partial pooling corrects allele proportion of Xist towards either of monoallelic expressions, but Xist is still expressed from both alleles in many cells. **(b)** Therefore, for many X chromosome genes, XCI is not completely locked in over the developmental stages of blastocyst cells. Partial pooling also corrects the ASE states and displays maternal allele preference more clearly for both unique reads and weighted allocation, although overall incompleteness of XCI is common irrespective of ASE estimation methods.

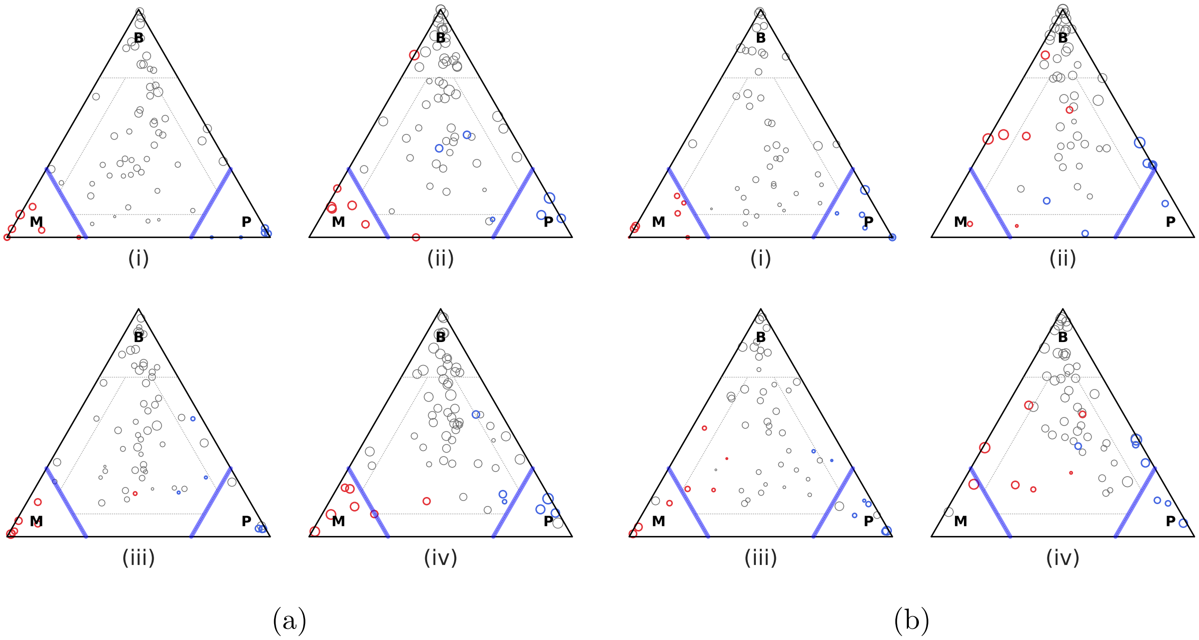


Supplemental Figure S12: Allelic expression state of putative imprinted genes for 113 and 125 primary mouse fibroblast cells[24] from **(a)** B6×CAST and **(b)** CAST×B6 hybrids respectively. We estimated the proportions of cells in paternal monoallelic, bi-allelic, maternal monoallelic states, $(\pi^P_{g\cdot}, \pi^B_{g\cdot}, \pi^M_{g\cdot})$, for imprinted genes based on (i) unique reads, (ii) weighted allocation, (iii) unique reads with partial pooling, and (iv) weighted allocation with partial pooling. Imprinting is not manifested for most of these genes. This observation holds across all methods of estimation including the unique-reads method. Some imprinted genes identified by unique-reads method (denoted in red and blue) are not in **M**- or **P**-class weighted allocation of multi-reads or partial pooling, for the reasons we discussed in Supplemental Figure S1. However, there are several clear examples of imprinting too. For example, Reinius et al. identified 13 from the 158 putative imprinted genes[26, 27, 28]. Using `scBASE`, we identified 14 imprinting genes (*B830012L14Rik*, *Gpr1*, ***Igf2r***, ***Impact***, *Mecp2*, ***Meg3***, ***Mirg***, *Ndn*, ***Peg3***, *Peg10*, ***Peg12***, ***Plagl1***, *Pon3*, ***Rian***) after weighted allocation and partial pooling – 8 of them (in bold text) are common to what Reinius et al. reported.

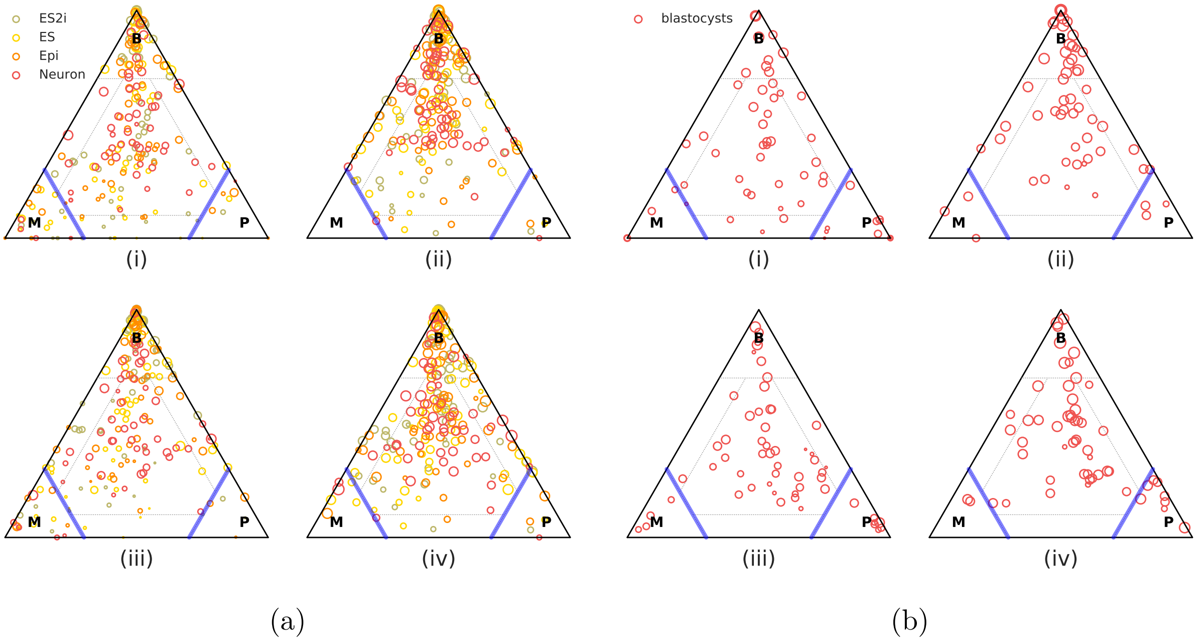


Supplemental Figure S13: Allelic expression state of putative imprinted genes **(a)** 231 mESCs, 183 mEpiSCs, and 74 motor neuron cells[25], and **(b)** 122 pre-implantation mouse blastocyst cells[7]. We estimated the proportions of cells in paternal monoallelic, bi-allelic, maternal monoallelic states, $(\pi^P_{g\cdot}, \pi^B_{g\cdot}, \pi^M_{g\cdot})$, for putative imprinted genes[26, 27, 28] based on (i) unique reads, (ii) weighted allocation, (iii) unique reads with partial pooling, and (iv) weighted allocation with partial pooling. As in fibroblast cells, imprinting is not manifested for most of the genes in these cells. This observation holds across all methods of estimation including the unique-reads method. Out of the 158 putative imprinted genes, `scBASE` identified 8 (*H19, Meg3, Nespas, Peg3, Snrpn, Tnfrsf22, Xlr3b, Zfp42*) and 10 (*Dhcr7, Impact, Lin28b, Peg3, Peg12, Ppp1r9a, Sfmbt2, Tnfrsf22, Xlr3b, Zrsr1*) genes after weighted allocation of multi-reads and partial pooling respectively.
